# Uncovering tissue-specific binding features from differential deep learning

**DOI:** 10.1101/606269

**Authors:** Mike Phuycharoen, Peyman Zarrineh, Laure Bridoux, Shilu Amin, Marta Losa, Ke Chen, Nicoletta Bobola, Magnus Rattray

## Abstract

**Motivation:** Transcription factors (TFs) can bind DNA in a cooperative manner, enabling a mutual increase in occupancy. Through this type of interaction, alternative binding sites can be preferentially bound in different tissues to regulate tissue-specific expression programmes. Recently, deep learning models have become state-of-the-art in various pattern analysis tasks, including applications in the field of genomics. We therefore investigate the application of convolutional neural network (CNN) models to the discovery of sequence features determining cooperative and differential TF binding across tissues.

**Results:** We analyse ChIP-seq data from MEIS, TFs which are broadly expressed across mouse branchial arches, and HOXA2, which is expressed in the second and more posterior branchial arches. By developing models predictive of MEIS differential binding in all three tissues we are able to accurately predict HOXA2 co-binding sites. We evaluate transfer-like and multitask approaches to regularising the high-dimensional classification task with a larger regression dataset, allowing for creation of deeper and more accurate models. We test the performance of perturbation and gradient-based attribution methods in identifying the HOXA2 sites from differential MEIS data. Our results show that deep regularised models significantly outperform shallow CNNs as well as k-mer methods in the discovery of tissue-specific sites bound *in vivo*.

**Availability:** For implementation and models please visit https://doi.org/10.5281/zenodo.2635463.

## 1 Introduction

Chromatin immunoprecipitation followed by sequencing (ChIP-seq) can reveal the genomic regions bound by transcription factor (TF) proteins in different tissues or developmental stages. To infer binding locations, short DNA reads are aligned to a reference genome assembly and peak calling techniques such as MACS [1] are used to localise the regions enriched in the IP experiment compared to a control. Inferred TF peak locations are typically hundreds to thousands of base-pairs in length and contain functional sequence motifs identifiable as highly over-represented short k-mers or position-specific score matrices (sequence motifs, usually 6-10nt), corresponding to the binding locations of regulatory TFs. Widely used motif discovery tools include MEME [2], Homer [3], GEM [4] and KSM [5]. These tools can be used to annotate and visualise over-represented motifs using databases of known TF binding sites.

TFs frequently cooperate to achieve their cell-type binding specificity. Binding in different tissues may be enhanced by the presence of specific co-factors [6]. Several modes of cooperation are possible, including heterodimer formation (direct) or changes in the affinity of neighbouring sites as a result of increasing chromatin accessibility (indirect). Therefore, differential binding of a major regulator in different cells can be highly informative about cell-type specific TF interactions. For example, the MEIS homeodomain TFs are major developmental regulators in vertebrates and cobind with a large set of other factors [7, 8]. MEIS bind to a large proportion of accessible chromatin in mouse branchial arch tissues and are essential for development of this embryonic region [8]. HOXA2 is expressed concurrently with MEIS in the second branchial arch (BA2) and posterior branchial arches (PBA), but not the first branchial arch (BA1) (see Fig. 1), and was shown to cooperatively bind with MEIS in BA2, resulting in a mutual increase of occupancy [8]. Based on these observations, we reasoned that differential analysis of MEIS binding could reflect co-binding with specific partners in the BAs, including developmentally important HOX TFs.

**Figure 1:**
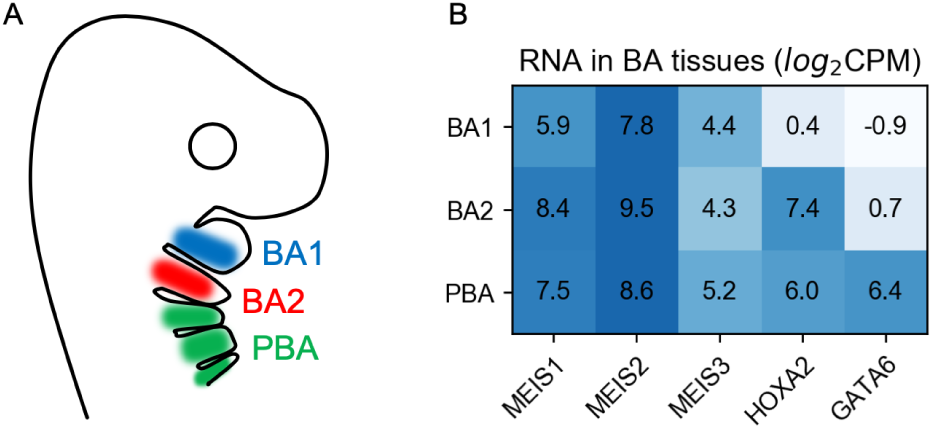
A) Location of BA tissues marked on a cartoon of mouse during embryonic development. B) Amount of RNA measured by RNA-seq in BA tissues. For ChIP-seq experiments a pan-MEIS antibody was used to immuno-precipitate MEIS 1-3. CPM - counts per million sequenced reads.

Deep learning approaches such as convolutional neural networks (CNNs) became state-of-the-art in visual and speech applications, followed by their application in genomics [9]. DeepBind [10] was the first model to use CNNs to identify DNA and RNA binding sites by training a network to classify between binding regions and a randomly shuffled negative set. DeepSEA [11] included epigenetic and accessibility data to jointly learn and predict the effects of sequence mutation (later expanded in ExPecto [12]), while FactorNET [13] extended the convolutional architecture with a bi-directional recurrent network to predict the ChIP-seq profile along the sequence. Recent works also include a GAN-based generative model for sequence [14], the *Basenji* network for prediction of gene expression [15], modelling binding from reporter assays [16], predicting differential expression from histone marks [17], and ensemble bootstrap models for handling imbalanced data [18].

Differential feature identification in genomic sequences can be accomplished in several ways. In k-mer approaches all possible combinations of nucleotides (up to a certain length) are counted in the differentially-bound regions and their frequencies compared with a background set. After enriched k-mers are identified (and possibly combined to a position-weight matrix, PWM), the sequences are scanned for alignment with the motif. Counting is increasingly time-consuming for longer k-mers, and annotation of the genome with a PWM is insensitive to the sequence features surrounding it. Deep learning models do not allow easy visualisation of features in general due to high non-linearity, but can attribute them in an input-dependent manner. This means that compared to a k-mer approach the same motif can be identified as a feature with different importance, depending on the context in which it appears in the region. The simplest 1-layer CNN is similar to a k-mer method in that it learns to identify regions based on the statistical occurrence of a number of PWMs, represented as convolutional filters. In a deep learning model these are optimised simultaneously with classification or regression parameters that follow. Deeper convolutional networks are able to learn spatial patterns with a wider receptive field, but require more training data in order to fit more parameters.

Prediction attribution refers to identifying the elements of the input which caused the neural network to predict a given output. *In silico mutagenesis* is a perturbation-based approach introduced with DeepBind, which uses the model to predict effects of all possible single-nucleotide substitutions in a region, creating a mutation map. This approach can be computationally expensive when predicting saturated mutation in larger regions or for more than one nucleotide at a time. Alternative approaches seek to approximate the Shapley value and satisfy the axiom of *completeness* [19], also known as summation-to-delta. This requires distributing the difference in model prediction between a reference and the input on the elements of the input. *Integrated gradients* and *DeepLift* [20] are two approaches that allow this. Because DeepLift distributes the activations in a model-specific manner we chose to evaluate integrated gradients, which are implementation independent. In this approach, gradients are calculated over a number of steps, while linearly interpolating between the example and a reference, finally multiplying by their difference. This captures the non-linearity of a deep model in the attribution. A reference is a background example, which ideally contains no features. All zeros can be used (in the case of one-hot encoded sequence data) which is conceptually similar to using a black image in a vision application. Multiplying *gradient times input* is a fast method of obtaining attribution, and a special case of integrated gradients with a reference of zeros and a single integration step. Specifying reference for a genomic sequence is problematic due to categorical encoding, as linear interpolation between two one-hot samples does not result in another one-hot sample. Similarly, prediction for an all-zero input is not well defined for a network trained using one-hot examples.

In a high-dimensional problem, model identifiability becomes an issue. Deep models with millions of parameters can be particularly difficult to train on smaller datasets because the loss landscape contains many local minima. As a result the attribution becomes unstable and initialisation-dependent. Typical methods of regularising the model include transfer learning [21], where a portion of neuron weights is transferred from a model trained on data from a related domain, and semi-supervised learning, where a large unlabelled dataset is used in a parallel training task. In our case, a large dataset with regression targets is available in several replicates, from which we also have a much smaller subset of confidently labelled differential examples.

In this contribution we extend the use of deep learning models to the identification of sequence features predicting differential TF binding. We use CNN models to identify DNA sequence features predictive of changes in ChIP-seq data across conditions. To illustrate the problem of differential binding, we use MEIS ChIP-seq data from BA1, BA2 and PBA in mouse embryos. It was previously shown that HOXA2 is the primary co-factor of MEIS in BA2 [8], which makes its experimental binding profile useful for validating attribution. For validation we train models to predict differential binding (relative increase or decrease in occupancy) from input sequence, and attribute the prediction to nucleotides in each region. HOXA2 binding sites positively contribute to MEIS occupancy in BA2 and PBA, and therefore appear as features of the BA1-downbinding class. Our approach is illustrated in Fig. 2, which shows an example where learned features predictive of differential MEIS binding are consistent with co-binding of HOXA2 and MEIS. We compare the accuracy of k-mer approaches to CNNs used with mutagenesis and integrated gradients, for which we validate performance with a zero background, as well as averaging 10 real genomic backgrounds. These are selected randomly from enhancer regions from H3K27ac ChIP-seq peaks with no detected MEIS binding. We then compare the locations of features ranked by a sliding window to the *in vivo* ChIP-seq profile of HOXA2 in BA2, in two experimental replicates. The HOXA2 data is not used in training the networks and therefore provides independent validation of the learned features. We create deep learning models using regression of all available replicate data in order to regularise the classification task, increase predictive performance, as well as accuracy and stability of feature attribution.

**Figure 2:**
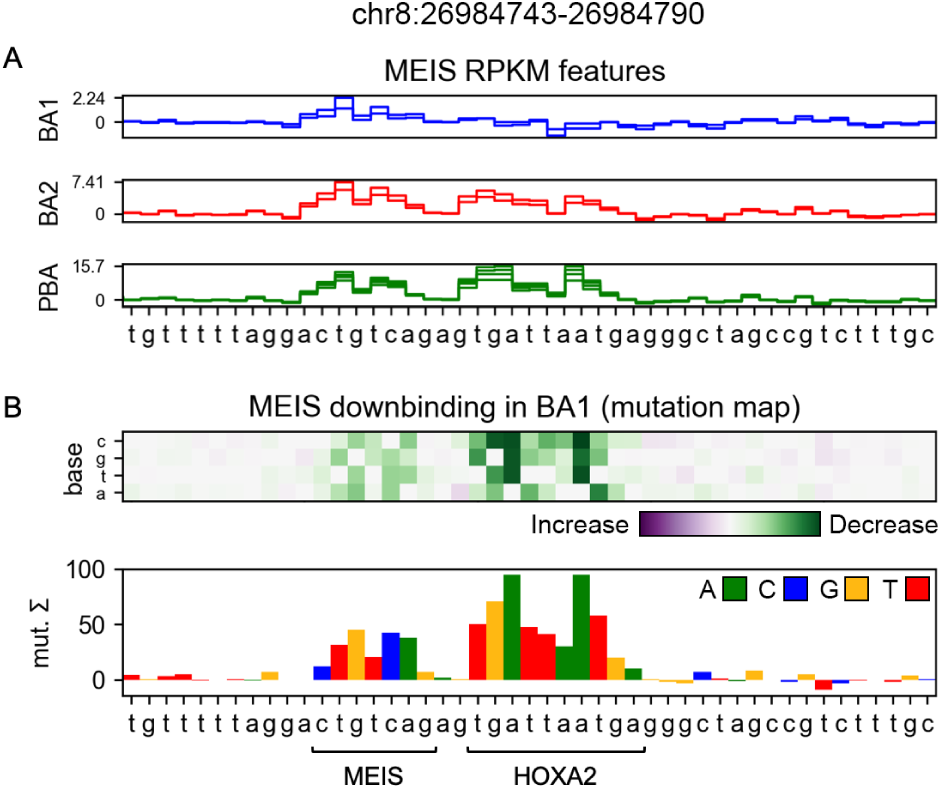
A) MEIS RPKM (reads per kilobase of transcript per million mapped reads) regression features attributed by a deep model using mutagenesis. Values indicate the sum of predicted RPKM change if the base was mutated to all of its alternatives. Each line indicates features of a single replicate output for a tissue. Colours match cartoon labelling in Fig. 1A. MEIS binding site (CTGTCAG) is a feature in all tissues. B) BA1-downbinding features from a differential model. A dimeric site containing HOXA2 and MEIS binding motifs is identified as a differential feature enhancing MEIS binding in BA2 and PBA, but not BA1.

## 2 Methods

### 2.1 Data accession and processing

To identify co-binding features of MEIS in the BA tissues of interest we obtained genome-wide binding profiles from MEIS ChIP-seq experiments. ChIP-seq results vary in quality, which motivates the use of several biological replicates. We used previously published data from ChIP-seq for MEIS, HOXA2 and H3K27ac [22, 8, 23] which we re-analysed for the mouse mm10 build. The ArrayExpress accession numbers for the data sets are: E-MTAB-7766, E-MTAB-7767, E-MTAB-5394, E-MTAB-5407, E-MTAB-5536, E-MTAB-7963, E-MTAB-7966. Pre-processing of the ChIP-seq data was identical to the original papers (Trimmomatic for trimming [24], Bowtie2 for aligning to the mouse genome [25], samtools [26] to remove the aligned reads with a mapping quality Q30 and MACS2 for peak calling [1]), followed by Diffbind [27] recentering to the position common across replicates. RPKM (reads per kilobase of transcript per million mapped reads) values are calculated for peaks, measuring the amount of binding. To identify differential occupancy, we use edgeR [28] with TMM normalisation. Labels are assigned to regions which show either increased or decreased level of binding in one tissue compared to the other two, and a non-differential label is given to regions without significant difference in RPKM across tissues. Label counts obtained this way are shown in table 1. For input to neural networks the sequences are one-hot encoded to a fixed-length 2-dimensional array *BxL*, where *B* = 4, representing each of the possible bases, and *L* is the chosen length of encoded sequences. At each base position the array is 1 for the present base, and 0 otherwise. In order to constrain computational cost the length is bound to between 200 and 2000 nucleotides. RNA-seq gene expression values used in this paper are identical to originally published [8, 23].

**Table 1:** Differential labelling of MEIS-bound regions.

| Type | Count | Avg. length (nt) |
| --- | --- | --- |
| Increased binding |  |  |
| BA1 | 3416 | 778.9 |
| BA2 | 3850 | 790.2 |
| PBA | 18088 | 770.1 |
| Decreased binding |  |  |
| BA1 | 2345 | 860.7 |
| BA2 | 3070 | 867.2 |
| PBA | 17923 | 857.7 |
| Non-differential | 127185 | 633.6 |
| All MEIS | 215830 | 679.3 |

### 2.2 K-mer-based methods

For k-mer attribution we used Homer [3] to identify enriched PWMs *de novo* by contrasting the regions in a differential class with the non-differential background. We then annotated the differential regions with most confident PWM, sorting locations from strongest to weakest match. While PWM is convenient for visualisation, the identified representation assumes independence between nucleotides. KSM (k-mer set memory, [5]) is an alternative method, which does not combine the individual k-mers into a single frequency matrix, but lists and ranks all identified instances independently. Likewise, we annotated the regions with the k-mers identified by KSM in order of confidence. Details of both approaches are given in the supplementary material.

Support Vector Machine (SVM) models split the input space with a hyper-plane boundary. During training optimal support vectors are found, which define the plane that maximises the separation of the training examples, while considering an adjustable soft margin for outliers. In order to allow classification of data which are not linearly-separable, a kernel function is used to calculate the distances between examples in a higher-dimensional space. In case of DNA sequences, the distance between two samples can be calculated based on differences in k-mer frequencies. Gapped k-mer SVM (GKM-SVM [29]) introduces the gkm-kernel, which allows for a given number of mismatches between the example sequence and the k-mers used as features (typically up to 4 mismatches for *k* = 11), and efficiently computes the distance based only on k-mers present in either sequence of each pair. We evaluate the more memory efficient implementation LS-GKM [30], which unlike the original does not pre-compute the entire distance matrix between all pairs of samples. Additionally, it introduces expansions of the gkm-kernel, which increase the contribution of k-mers found in the centres of peaks (wgkm-kernel), and alongside it apply a radial basis function to the identified k-mer frequencies (wgkmrbf-kernel). We compare the predictive and attribution performance of SVM to CNN models. Details of training and attribution with mutagenesis and GkmExplain, an integrated gradient method [31], can be found in the supplementary material.

### 2.3 Deep learning models

#### 2.3.1 1-layer CNN

For our baseline model with one convolutional layer we use an extended version of DeepBind [32], in which the convolution is followed by global *max*-pooling and parallel *average*-pooling, outputs of which are concatenated before being passed to fully-interconnected layers performing classification or regression. This CNN is capable of recognising spatial dependencies between nucleotides up to the length of its convolutional filters. Combinations of motifs in a region can still be recognised beyond that length through the pooling statistics, but their mutual distance is invariant to the network. Using *max* as the only pooling operation works well for classification, but manifests an issue in perturbation-based attribution if more than one motif of the same kind is present in the region. A concatenation of two types of pooling seems to alleviate this problem, and works well for classification as well as regression.

#### 2.3.2 Deep CNN

Our deeper models are based on the architecture of Basenji [15], additionally expanded with bottleneck layers (see Fig. 3A). In the initial layers of the network we instantiate repeating blocks of convolution, batch normalisation, 1x convolution (bottleneck), dropout and *max*-pooling. These pooling blocks reduce the spatial dimensionality of the input. The bottleneck layer was shown to improve performance of the DeepSEA network with linear projection [33]. In computer vision applications this kind of layer (a 1×1 convolution for 2D images) often uses a non-linear activation function. We validated the performance of models with and without the bottleneck, and with linear or ReLU (Rectified Linear Unit) activation. The second type of block uses dilated convolution instead of pooling to further expand the receptive field, while maintaining a constant output width. The dilation blocks are concatenated to form a hyper-residual network [34, 35]. The output is obtained from a linear 1x convolution and global average pooling. Unlike Basenji which used a Poisson loss, we perform our regression with a mean squared-error (MSE) loss on log of RPKM values.

**Figure 3:**
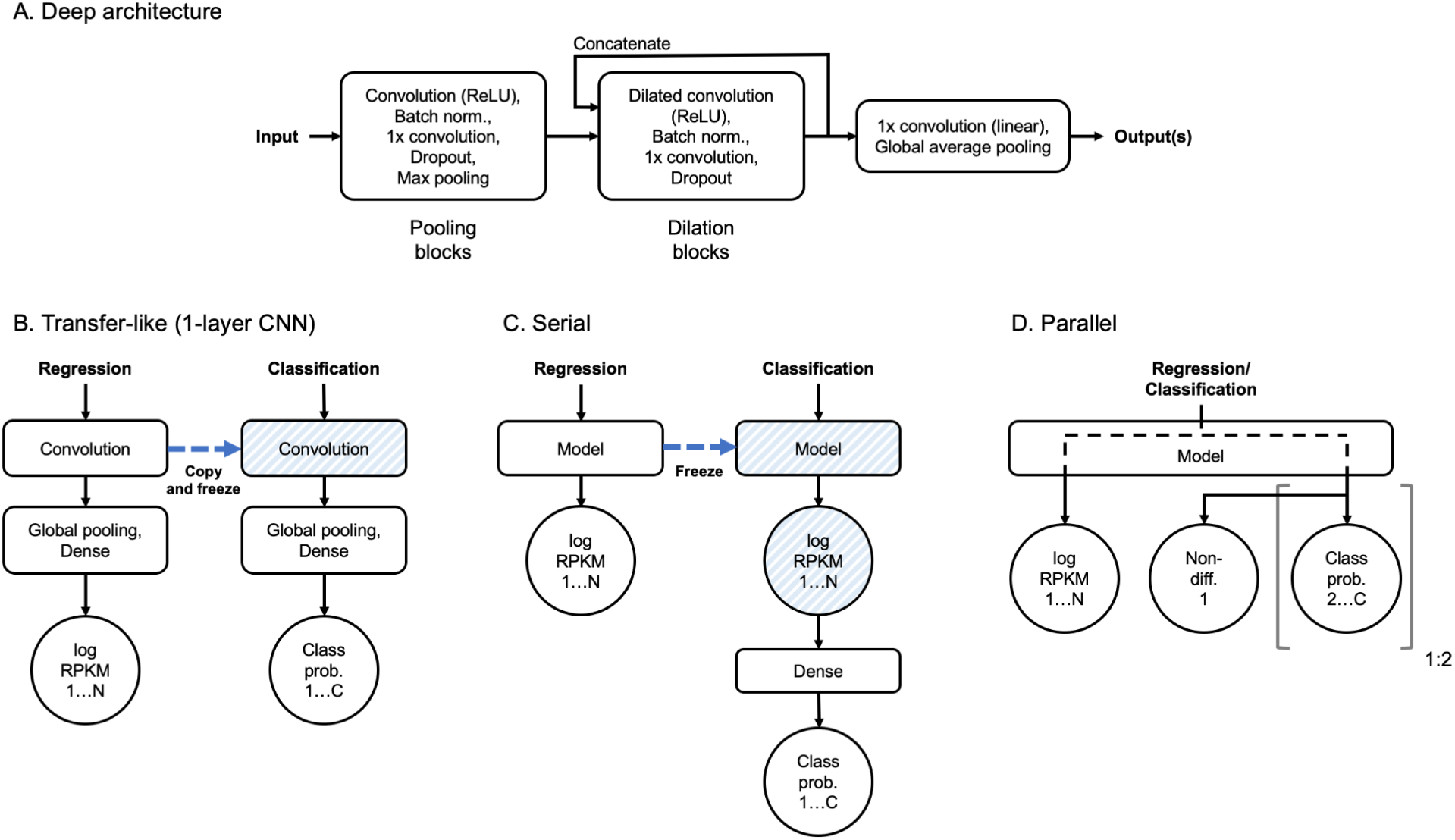
A) Schematic of a deep architecture. Pooling and dilation blocks are repeated for desired number of times, dilation blocks having their outputs concatenated. Number of blocks and other hyper-parameters are automatically optimised in the process of model selection. Input is a one-hot encoded nucleotide sequence. One or more outputs can be specified for classification and/or regression with task-dependent activations and loss functions. B-D) Modes of regularisation where latent variables of a larger dataset (regression, N targets) are used to regularise the training with a smaller dataset (classification, C classes). B) Typical use of transfer learning in 1-layer CNN. A convolutional layer is copied and frozen for training of the second model, allowing for inference in terms of previously learned intermediate latent variables. C) Serial architecture uses the output of the trained model as its input, performing non-linear weighting of regression targets for classification. D) Parallel architecture alternates between training model outputs in each batch. In this architecture latent variables are shared throughout model depth. Non-differential target is shared between up-binding and down-binding task outputs if both are used.

### 2.4 Model selection and training

For model selection we used the *Adam* optimisation method [36] and random search (see supplementary Fig. S1). Hyper-parameter ranges and model-specific settings are detailed in supplementary table S1. In each case a fifth of the data is held out for test and 3-fold cross-validation with early stopping is performed on the remaining part. The mean of training losses is calculated at the stoppage points and subsequently used as a stopping criterion when the final model is trained on the entire cross-validation data. For all models the input length is a hyper-parameter between 200 and 2000 nucleotides, which for a 1-layer CNN it is sampled at random. For the deeper models the receptive field (the maximum span in the input which affects activation prior to the global pooling layer) is calculated (supplementary algorithm 1), and randomly expanded up to twice to obtain the input length. Input length of the base RPKM models determines this length for subsequent classification for both transfer-like and serial approaches. In the training and validation sets we augment the least frequent down-binding classes with reverse complement sequences. Augmentation is performed on all BA1-down examples, and on the remaining classes only if their label count is below the augmented BA1-down.

### 2.5 Input-level attribution with neural networks

In-silico mutagenesis is performed using methods defined in DeepBind [32], which we adapted for variable-sized regions and model inputs (see supplementary for details). For each region of interest a mutation map of all single-base substitutions is created. From this map, individual importance of each nucleotide is calculated by summing change in model prediction caused by mutation to the alternative bases. For integrated gradients we evaluate using a zeros reference, as well as averaging the attribution with 10 enhancer regions as references, obtained from non-differentially classified H3K27ac peaks without detected MEIS binding. We determine the number of integration steps by calculating summation-to-delta over a number of regions (supplementary Fig. S6). For all methods we use raw logit activations preceding the final *softmax* layer. After obtaining attribution, a sliding window approach is used to identify the locations of strongest features in the dataset, and rank them from strongest to weakest. In this application the attribution is performed on the same dataset used for training. Although generalisation and overfitting are usually considered in terms of predictive loss, not feature attribution, to ensure generalised features we additionally train models holding out every fold of the data, and use them to attribute over the held-out folds. We validate attribution using each fold model individually, an ensemble of models, and a single model trained on all the data.

### 2.6 Regularising high-dimensional problems with multitask learning

For a baseline we train a DeepBind-like 1-layer CNN directly on the classification dataset. To regularise this architecture, we adopt a transfer-like methodology, in which a regression model is trained to predict RPKM values for all tissues and replicates, and the convolutional layer is copied to a new model and frozen (disabling gradient descent updates during training, see Fig. 3B). Model selection is subsequently performed for the classifier parameters that follow the convolution. Similarly, we perform model selection for a reference deep CNN using only the classification dataset. We expect this model to overfit and be unstable due to being heavily over-parameterised, and therefore adopt two regularisation approaches. In the serial approach (Fig. 3C) a deep RPKM regression model is trained first and frozen, mimicking the re-use of a convolutional layer in a shallow CNN. The output log-RPKM values are used as the input to a shallow classification network. This allows for data-driven learning of the appropriate replicate weighting based on labelling of classification regions. Importantly, this approach exploits the sequential nature of the classification labels originating from the regression values. Secondly, we create parallel models (Fig. 3D) which contain task-specific training paths jointly spanning most of network depth, finally diverging to separate regression and classification outputs. Two classification tasks (up-binding and down-binding) can be specified with a shared non-differential class. In this case all of the shared parameters are updated when the paths are alternatively trained in batches, with the auxiliary regression loss not being part of the early stopping criterion.

### 2.7 Evaluation

#### 2.7.1 Model performance and attribution stability

The performance of regression models is evaluated by Pearson (R) and Spearman (Rho) correlation coefficient between each replicate target prediction and ground truth on the held-out test set. Between-replicate correlation within the same tissue is reported as an expected upper bound of model performance (see table 2). Classification performance is evaluated by precision-recall curves for each class and area under the curve (PR-AUC). Additionally, confusion matrices, per-class recall, and average class F1 score is calculated for each model. For feature identification we prioritise recall over precision, due to conservative labelling of differential regions which increases the chance of real weakly differential examples to be assigned non-differential ground-truth labels. This is expected to lower precision, despite the models identifying correct features. To measure stability of attribution to model initialisation we train 10 models (using the same data and hyper-parameters), and use them to attribute over randomly selected 1000 BA1-down regions. For each region a single feature is selected (see supplementary Fig. S7), and 25nt binary mask is created over this feature. The binarised attribution is compared between model instances by feature stability estimator [37] resulting in a score between 0 (random features) and 1 (identical features), which is averaged for all tested regions.

**Table 2:** MEIS cross-replicate and regression test correlation.

| Tissue | Replicate | Cross-replicate |  | 1-1 CNN |  | Deep CNN |  |
| --- | --- | --- | --- | --- | --- | --- | --- |
|  |  | R | Rho | R | Rho | R | Rho |
| BA1 | 1 | 0.605 | 0.634 | 0.43 | 0.39 | 0.51 | 0.46 |
| BA1 | 2 | 0.605 | 0.634 | 0.34 | 0.32 | 0.4 | 0.38 |
| BA2 | 1 | 0.685 | 0.690 | 0.42 | 0.42 | 0.47 | 0.47 |
| BA2 | 2 | 0.685 | 0.690 | 0.44 | 0.42 | 0.52 | 0.5 |
| PBA | 1 | 0.644 | 0.652 | 0.44 | 0.42 | 0.52 | 0.51 |
| PBA | 2 | 0.710 | 0.728 | 0.53 | 0.52 | 0.59 | 0.59 |
| PBA | 3 | 0.683 | 0.704 | 0.52 | 0.51 | 0.57 | 0.58 |
| PBA | 4 | 0.708 | 0.725 | 0.54 | 0.54 | 0.6 | 0.61 |

#### 2.7.2 Motif-centre Poisson test with ChIP-seq

Feature attributions are obtained for the BA1-down class from deep models, or k-mer counting annotations, and compared to two HOXA2 ChIP-seq replicates in BA2. HOXA2 is the dominant co-factor of MEIS in this tissue [8], which allows for direct validation of this class feature. To evaluate the identified locations, we select the strongest feature in each region and test against a background assuming Poisson distribution of reads, similarly to MACS2 [1], using a 500nt window around the feature, as in GEM [4] (see supplementary for details). In order for a feature to pass the test a p value <0.05 is required for peak alignment with both of the reference replicates. For each method we sort the features by method-specific score from strongest to weakest and report the proportion of features passing the test as the number of included locations is increased.

## 3 Results

### 3.1 Regularisation with a large dataset allows training of deeper and more stable models

By using regression data for model regularisation we were able to train deeper, highly-parameterised models with a wider receptive field, and obtain higher regression (table 2) and classification performance (Fig. 4, supplementary table S2 and S3) compared to shallow CNNs. Deep models also show increased attribution stability despite larger model size (table 3). Training a deep model without this regularisation results in poor predictive test performance and less stable attribution. The addition of the up-binding task in the 3-task parallel model increased overall predictive performance and reduced the number of parameters compared to the 2-task, but decreased the accuracy of BA1-down attribution. During model selection we varied the number of dilation blocks and input size for each type of model. We observed peak performance in predicting MEIS RPKM using 3 pooling blocks and 7 dilation blocks, totalling 21 convolutional layers and over 4 million parameters. Our best models include bottleneck layers with a strong (x0.25-x0.5) dimensionality reduction and ReLU activation (Fig. 5).

**Table 3:** Attribution stability (BA1-down, 1000 regions, 10 models).

| Type | Input nt | N. params | Grad.*Input | Integ. (16, z.) | Mutagenesis |
| --- | --- | --- | --- | --- | --- |
| 1-l CNN | 200 | 335,652 | 0.29 | 0.17 | 0.29 |
| 1-l CNN | 600 | 34,916 | 0.6 | 0.63 | 0.56 |
| 1-l CNN | 2000 | 283,716 | 0.5 | 0.24 | 0.47 |
| 1-l Transfer | 600 | 106,564 | 0.59 | 0.50 | 0.54 |
| Deep, direct | 1100 | 9,374,980 | 0.63 | 0.61 | 0.58 |
| Deep, par., 2-t | 2000 | 8,128,996 | 0.65 | 0.72 | 0.69 |
| Deep, par., 3-t | 800 | 2,119,180 | 0.5 | 0.72 | 0.68 |
| Deep, serial | 1000 | 4,346,908 | 0.72 | 0.8 | 0.74 |

**Figure 4:**
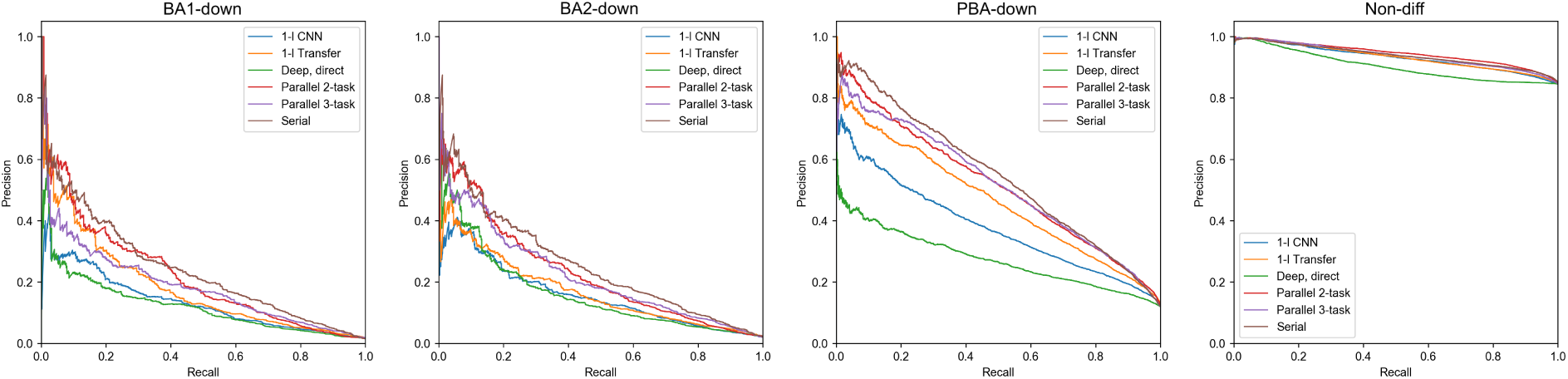
Test set precision-recall curves for the down-binding task. 1-layer CNN and deep, direct models were trained with classification dataset only. Transfer, parallel and serial models used MEIS regression data for regularisation.

**Figure 5:**
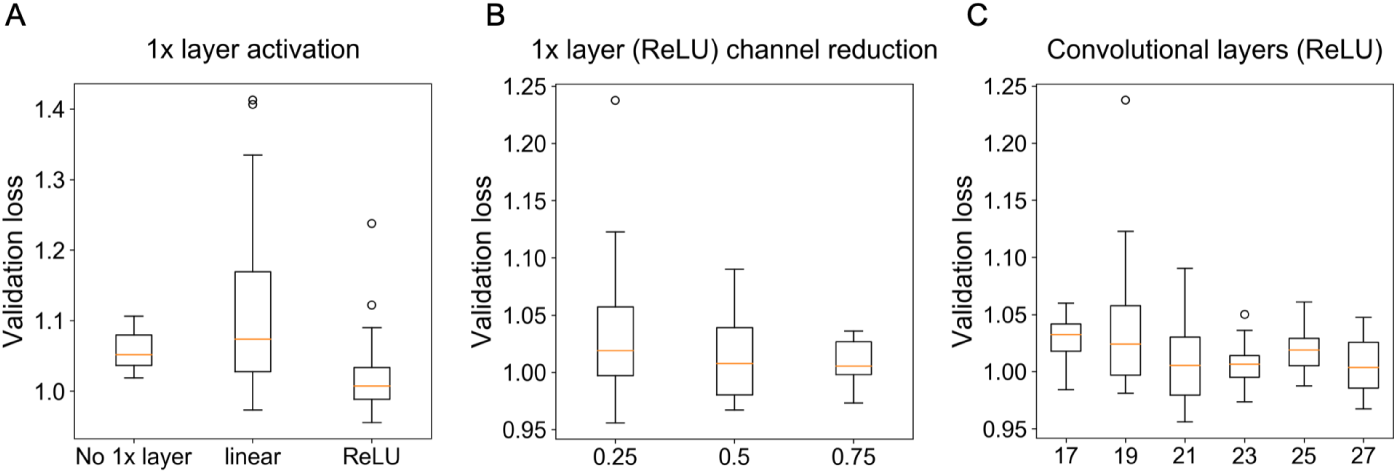
A) Validation loss of MEIS RPKM model selection when the 1x bottleneck layer is omitted, or used with linear or ReLU activation. B) Validation loss for varying amount of dimensionality reduction (proportion of channels of preceding layer) caused by the bottleneck using ReLU activation. C) Validation loss as a function of total number of convolutional layers (including 1x with ReLU activation) for MEIS RPKM regression model.

During model selection input length was automatically optimised and the resulting models differ markedly in the number of parameters. In particular, we observe that a 1-layer CNN becomes significantly over-parameterised when trained on a sub-optimal input length. The optimal 1-layer model with 600nt input has 9.6 times fewer parameters to best performing model for a smaller 200nt input, and 8.1 times fewer than model with 2000nt input, resulting in more stable attribution (see table 3). The likely reason for this behaviour is the 200nt regions containing fewer real features, and model overfitting to noise as a result. We also tested performance of models with and without the reverse complement (RC) augmentation of the least frequent BA1-down class, observing significant increase in performance of the 1-layer CNN (see supplementary Fig. S2 and table S4). The increase in predictive accuracy does not necessarily appear in the augmented class, but rather in averaged F1 performance for all classes. The benefit of RC augmentation is smaller for deeper models, which due to increased non-linearity appear to generalise well to RC sequences.

### 3.2 Deep models significantly outperform shallow CNNs and k-mer counting in identifying HOXA2 bound sites

Our architectures allow for identification of HOXA2 sites bound *in vivo* with significantly higher precision than previously possible with k-mer methods, as shown in Fig. 6A. True HOXA2 sites are identified with higher accuracy than Homer, even if the latter is allowed to see the ground-truth data for counting (Homer *known*). Visualisation in Fig. 7 reveals example HOXA2 co-binding features discovered by differential region classification based solely on the MEIS ChIP-seq data. The models allow to identify sequence features of any of the predicted classes, therefore a 3-task model can be used to identify binding features of relative increased or decreased MEIS binding. Up-binding attribution of MEIS in PBA can uncover several types of features, as many TFs cooperate in this region. Among those, GATA is a known differential co-factor of MEIS in PBA, and can be discovered as feature of PBA-upbinding, as shown in Fig. 8. Additional features of this class are shown in supplementary Fig. S13 and S14. The 2-task parallel model (trained only for the down-binding and regularised with regression) performs best in attribution of the confidently labelled BA1-down regions. Transfer of regression parameters in 1-layer CNN improves attribution performance compared to training using only classification labels, but does not match the performance of deeper models. Feature accuracy of KSM and Homer used *de novo* is comparable to 1-layer CNN in BA1-down regions. KSM outperforms Homer for the most confident features, but shows lower accuracy in a broader set of regions. Weaker performance of KSM in our application is likely due to our method of input annotation using ranked k-mer matches. The results suggest that in this case a PWM can capture more context useful for ranking than an ordered list of discrete k-mers. The comparison of CNN models with gapped k-mer SVM on the BA1-downbinding task indicates that CNNs outperform SVM in predictive performance of binary classification (see Fig. S9), as well as attribution performance in predicting HOXA2 (see Fig. S10). Example features obtained from SVM are shown in Fig. S11. CNNs benefit from GPU acceleration in training and attribution (see table S5 and S6) and are therefore faster to create and evaluate given appropriate hardware.

**Figure 6:**
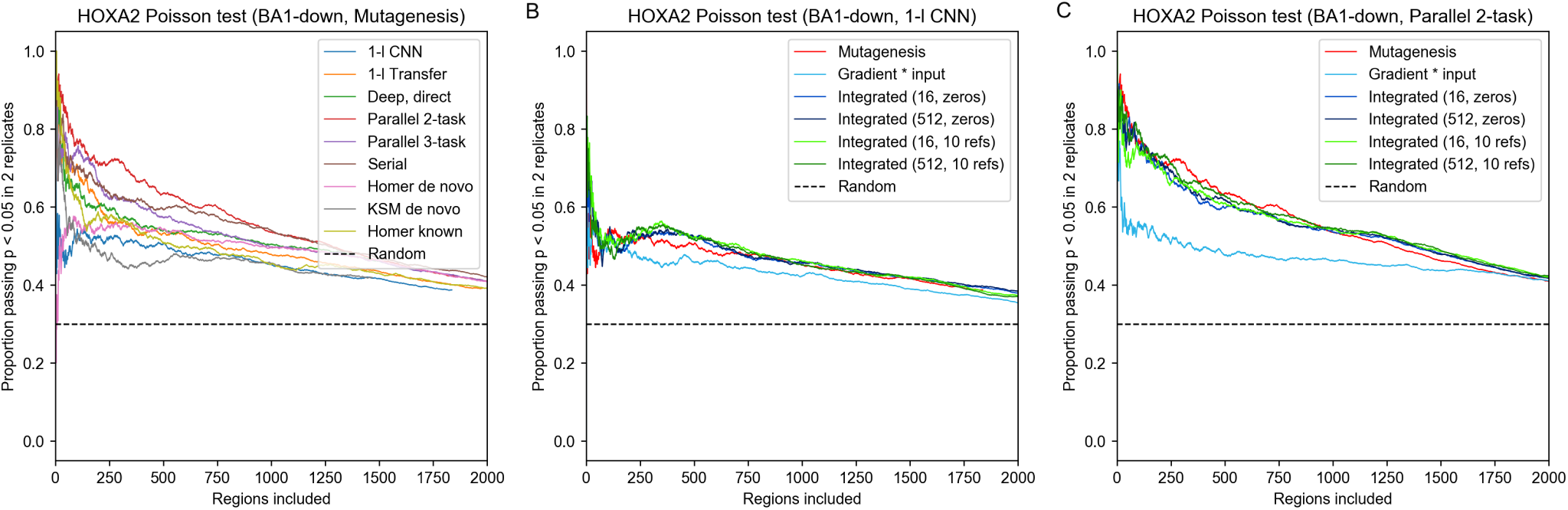
Proportion of most confident features identified by differential analysis passing a Poisson test for alignment with both HOXA2 ChIP-seq replicates. Regions labelled as BA1-down are tested. One strongest feature in each region is selected. Random indicates chance of randomly selected location in the regions passing the Poisson test. A) Comparison of CNNs with k-mer counting. Mutagenesis is used with CNN models. Homer known indicates using Homer with published HOXA2 ChIP-seq data (shown for reference). B) Attribution method comparison using 1-layer CNN. C) Attribution method comparison using deep parallel 2-task model.

**Figure 7:**
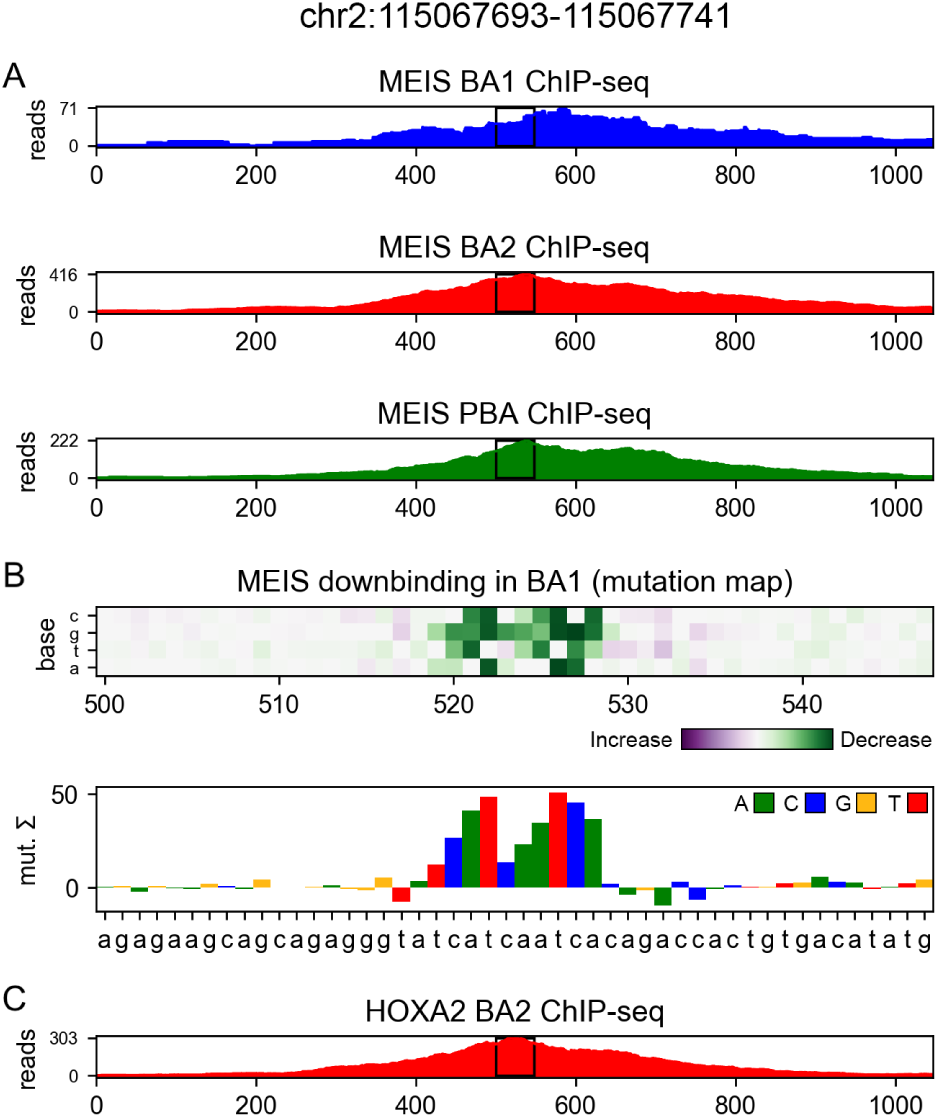
A) MEIS ChIP-seq profiles in a region differentially down-bound in BA1 compared to BA2 and PBA. B) Nucleotide-level mutation map (and its 1-dimensional channel sum), shown in the central region marked with black rectangles. Attribution of MEIS BA1-down differential class using 2-task parallel model identifies HOXA2 binding site (ATCAATC). C) Reference HOXA2 ChIP-seq profile (not used for model training).

**Figure 8:**
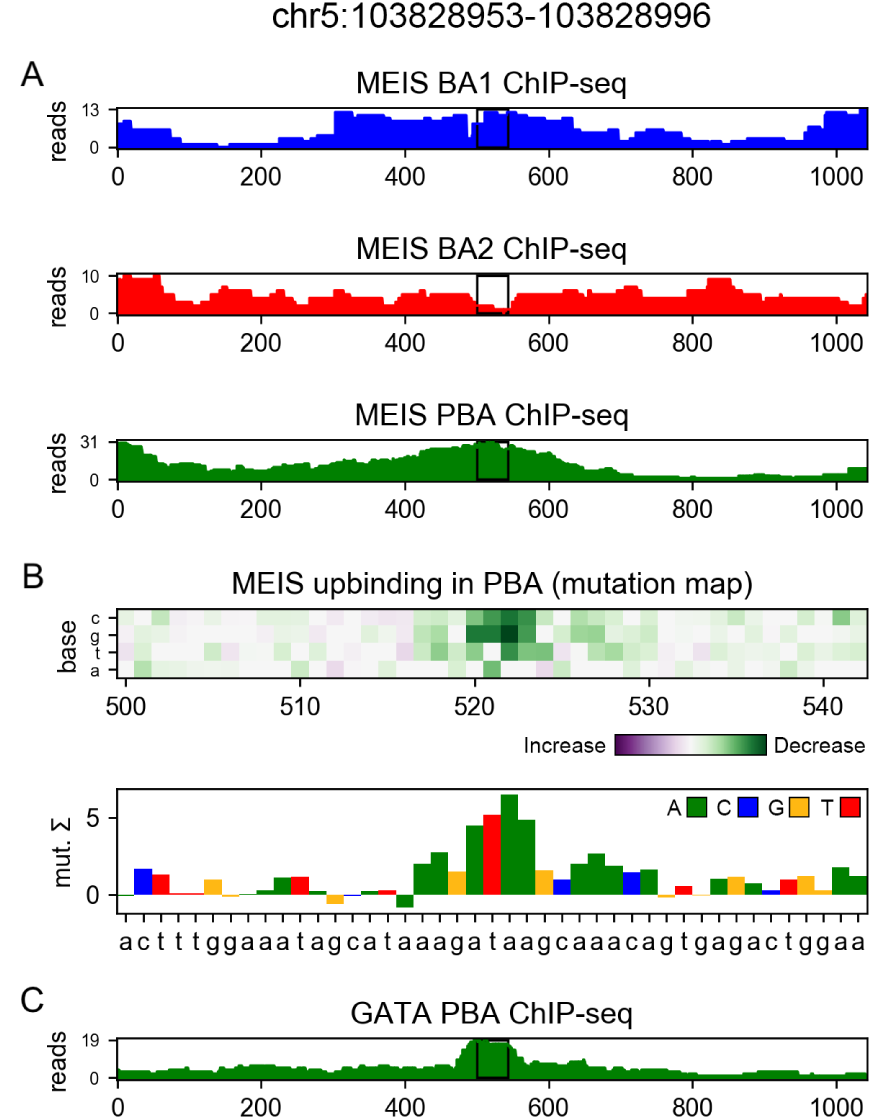
A) MEIS ChIP-seq profiles in a region differentially up-bound in PBA compared to BA1 and BA2. B) GATA binding site (AGATAAG) is identified as a feature of differential MEIS up-binding in PBA. Attribution was performed using mutagenesis and 3-task parallel model, and shown in the central region marked with black rectangles. C) Reference GATA ChIP-seq profile in PBA (not used for model training).

### 3.3 Differential analysis of MEF2D

In order to demonstrate the applicability of our approaches on another dataset, differential analysis of MEF2D [38] was performed across three mouse tissues (see table S8), and the results are shown in the supplementary. Transfer-like 1-layer CNN, as well as deeper serial models were created to regularise up- and down-binding classification tasks with regression data. The models show significant improvements over non-regularised CNNs, and deeper models provide increase in predictive performance over shallower ones (see table S7, S9 and S10, Fig. S15 and S16). Example regions of MEF2D co-binding with known TFs such as CRX [38] and MYOD [39] were identified (see Fig. S17), as well as other factors, some similar to previously reported as MEF2 interacting partners in other systems (see Fig. S18).

### 3.4 Mutagenesis performs similarly to integrated gradients in nucleotide-level attribution

We observe on our dataset that mutagenesis (using a scoring function from DeepBind, [32]) performs better or similarly well to integrated gradients in attribution accuracy (see Fig. 6 and supplementary Fig. S4), particularly with deeper models. Integrated gradients result in marginally higher attribution stability (excluding sub-optimal 200nt and 2000nt 1-layer models, see table 3). When specifying a background reference, 10 real regions consistently outperform a single all-zero reference. While our tests indicate that for the sum of attribution to reliably equal the difference in prediction (to within 5%) requires using as many as 512 integration steps (see supplementary Fig. S6), we observe that 16 steps perform nearly equally well for predicting HOXA2 binding, despite providing over-complete attribution. In this case exact completeness does not seem to be necessary for prioritisation of features. Gradient times input is generally outperformed by more computationally intensive methods, except for the 1-layer transfer-like CNN. We infer that as the depth and non-linearity of models increases, the gradient obtained at a single step is a poor predictor of model response to input perturbation. A significant increase in performance is observed when obtaining scores before the final softmax (similarly to [40]) for both integrated gradients and mutagenesis (see supplementary Fig. S3).

Performance of attribution can be further increased by training several models on different folds of the data, and averaging their attribution, as shown in Fig. 9. The benefit is evident in case of 1-layer CNN trained without regression data, and becomes smaller for transfer-like 1-layer CNN and deeper models, which are more stable across folds (see supplementary Fig. S8). Attribution with models which held-out the regions during training enforces generalised features, which appears to lower performance. The results suggest that holding out data may be detrimental to full attribution, especially for shallower models which are less able to generalise.

**Figure 9:**
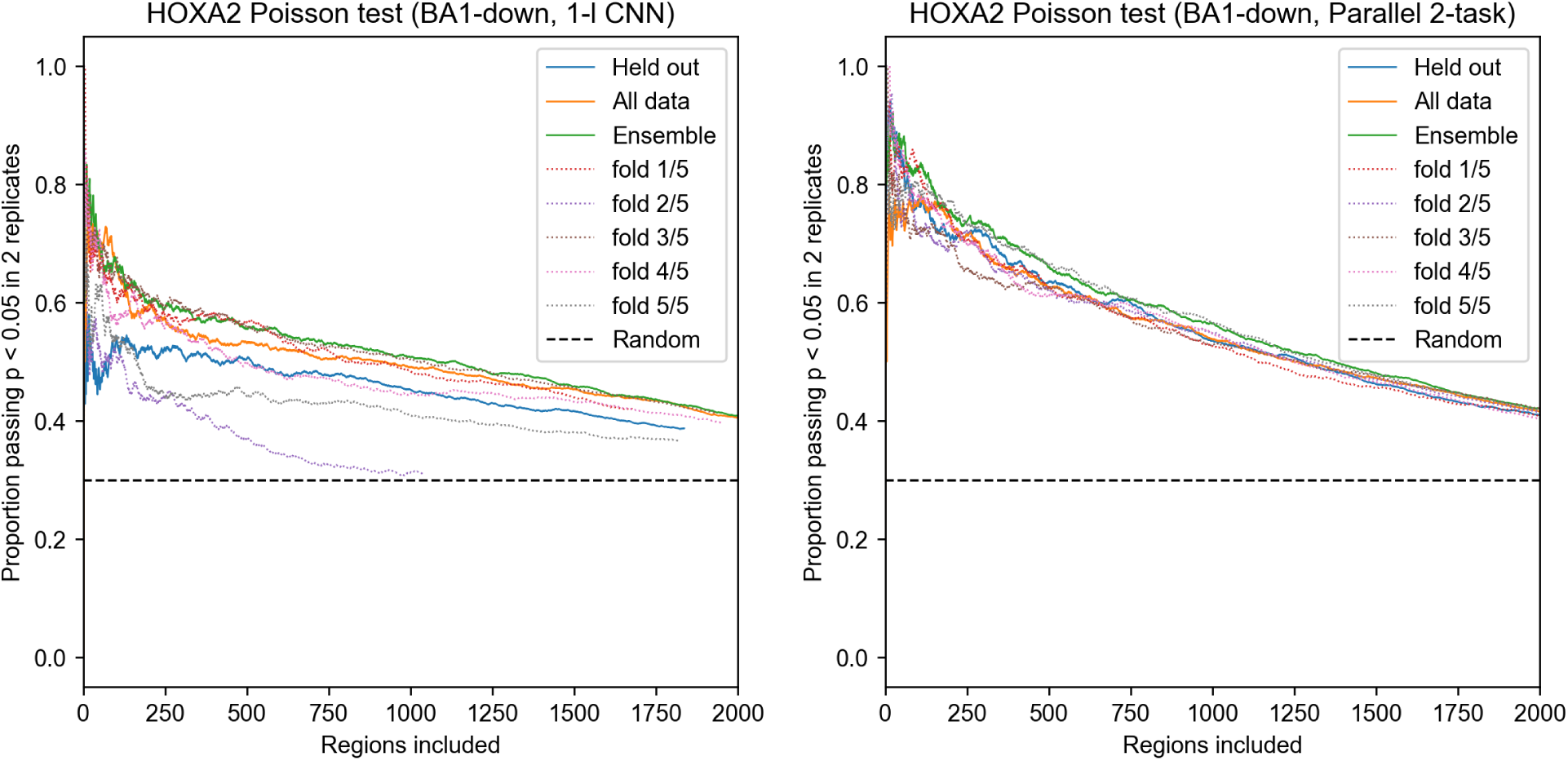
Overfitting effects in BA1-down mutagenesis attribution validated with HOXA2 ChIP-seq (Poisson test, p<0.05 in two ChIP replicates). 5 models were trained holding out different folds of randomly shuffled data. Held out indicates each peak was attributed with model which held out the region during training. Ensemble indicates using mean attribution from all models. All data indicates using a single model, trained on all the data.

## 4 Discussion

In this work we introduced CNN methods for identification of DNA sequence features predicting differential and cooperative TF binding. Using MEIS ChIP-seq data in mouse BA tissues we identified the binding locations of tissue-specific co-binding partners through differential classification of MEIS-bound regions. Validation with HOXA2 ChIP-seq showed that CNN models trained on MEIS data could reliably identify HOXA2 features in BA2, consistent with a synergistic effect of HOXA2 and MEIS binding [8]. Our results indicate that deep learning offers significant advantages over k-mer methods in identifying functional features *in vivo*, due to improved recognition of the context in which the motif appears in the region. This manifests particularly when attributing a wider set of regions, less confident *a priori* (see supplementary Fig. S5). Deep models lower the chances of false-positive attribution, and outperform Homer even if the latter is allowed to see the HOXA2 ChIP-seq used as the ground-truth. While our neural networks are able to recognise true binding sites with higher accuracy, k-mer methods remain useful in our workflow for clustering and annotating the resulting short features with known TF families. In a parallel work [41], cooperative binding properties of TFs are explored based on regression of ChIP-nexus profiles. Methods described here are distinct in that a differential objective function between cell-types is explicitly defined.

Training deep models on a relatively small and imbalanced classification dataset required using a larger set of regions for regularisation. While training parallel models for a specific task provides accuracy benefit, it is also time consuming (see supplementary Fig. S12). The addition of the up-binding task to the parallel model lowered the accuracy of the validated BA1-down attribution, despite increasing overall predictive performance. This is likely due to a decreased contribution of BA1-down class to the optimised loss. We observe that the serial model provides good attribution accuracy and stability, with the additional advantage of low training cost for new classes, as long as they can be predicted from regression targets. Inclusion of bottleneck layers with ReLU activation works well in our application. Since the hyper-parameter ranges used in model selection constrain the receptive field below the maximum allowed input size (2000nt), ReLU appears to provide a benefit in increasing the non-linearity of the model without increasing the receptive field. We achieve the best results with the highest tested dilation rate (4), suggesting a further increase of this parameter may be beneficial, especially for wider inputs.

Through evaluation of neural network attribution methods, we observe that single-nucleotide saturated mutagenesis performs well, and similarly to integrated gradients on our dataset. This appears consistent with good performance of mono-nucleotide models (such as [42]) indicating that single-nucleotide perturbations have a strong effect on binding.

In our opinion, approaches satisfying *completeness*, including integrated gradients and DeepLift, are particularly promising in domains where perturbation is less feasible (when operating on real-valued input), and where background samples can be easily specified. While higher-order saturated mutagenesis becomes computationally infeasible, not all combinations of substitutions are likely to be important in a given region. We note, however, that perturbation-based attribution is prone to adversarial effects and requires models trained on a large enough datasets in order to generalise well to unseen mutation. Our work shows that using deep learning, which increases non-linearity and provides a wider input context to a model, is beneficial in uncovering sequence features contributing to tissue-specific transcriptional regulation.

## Acknowledgements

We thank the members of the Genomic Technologies Facility, Ian Donaldson and the Bioinformatics Core Facility at the University of Manchester for processing the ChIP-seq data and Munazah Andrabi for supplying the expression data.

