## Supplementary material for "Uncovering tissue-specific binding features from differential deep learning"

### 5 Supplementary material

#### 5.1 Model selection and training

Keras 2.2.4 and Tensorflow (GPU) 1.12.0 are used for training, with Adam optimiser [1] (default settings). Hyper-parameter ranges are given in table S1. The 1-layer CNN is created by instantiating a 1-dimensional convolutional layer (implicitly spanning channel depth $= 4$) with given number of filters and filter length, ReLU activation and stride $= 1$. The output of this layer is globally pooled by separate $average$ and $max$ operations, and concatenated. Following layers consist of batch normalisation, dropout, and an optional dense layer with ReLU activation. Finally, model outputs are defined with *softmax* (classification) or linear (regression) activations.

The deep models are constructed by instantiating a given number of pooling blocks, followed by dilation blocks, followed by a final convolutional layer with task-specific outputs. The layers of a given block type share the same hyper-parameters (are of the same size). Three pooling blocks with a pooling rate of 2, and between 4 and 9 dilation blocks were used for the tasks in this paper, however, number of blocks in particular can be lowered or increased for other datasets. The two types of blocks use different approaches to receptive field expansion. Max-pooling is used initially to reduce dimensionality and therefore lower memory requirements. Residual blocks are added next, which connect the outputs of each previous block to inputs of all the blocks that follow. They use dilated (atrous) convolution in order to preserve width and allow concatenation along the channel dimension. Both types of blocks use bottleneck layers of length-1 convolution (ReLU), with batch normalisation before, and dropout after the bottleneck.

For training of transfer-like models, a 1-convolutional layer CNN is trained first for the regression task, and the convolutional filter parameters are subsequently reused in the classification CNNs. Similarly, a serial model is constructed by training a multilayer perceptron that follows the regression model. In both cases, the transferred parameters are not updated for classification to prevent overfitting. Parallel models created here used the deep model settings, however the concept of parallel training could be applied to any architecture. The input to the model is shared across all tasks, while the outputs are separate. Training is performed in batches by alternating task-specific outputs. Different batch sizes were used for regression and classification tasks, as well as a scaling factor for the classification learning rate (*LRF*). LRF divides the regression learning rate (0.001) and its range should be adjusted to approximate the proportion of regression to classification data. Sum of classification task losses is used as a stopping criterion. Regression loss is not used for early stopping.

Genomic regions are resized to desired model input width (constrained between 200nt and 2000nt and rounded to 100nt, randomly sampled for 1-layer CNN; for deep models, receptive field is calculated and multiplied by *expand RF* hyper-parameter), one-hot encoded, and shuffled. Classification tasks are optimised using categorical cross-entropy loss after applying softmax activation in the final layer. Regression is optimised using mean-squared error loss, on $log_2$ of RPKM counts. In order to report test performance a fifth of the data is held-out. The remaining part is used for 3-fold cross-validation. During the cross-validation hyper-parameters are uniformly sampled at random from the specified intervals. Models are trained on each of the 3 folds until the validation loss ceases to improve for a specified number of epochs (early stop). Up-sampling is used to handle imbalanced data (sampling the same number of examples at random from each class). Training loss is recorded at epoch of lowest validation loss. Both training and validation losses are averaged between the 3 folds. After a given number of hyper-parameter sets is sampled, the set with the lowest validation loss is selected. Model is then trained on the entire cross-validation data until the corresponding mean training loss is reached. Test performance of this model is reported on the held-out set.

Number of CV iterations is given in table S1 and is equal for all deep models, and larger for 1-layer CNNs due to shorter training time. 50 CV iterations were used for 1-layer CNN RPKM model. For validating the bottleneck layer (box plots in Fig. 5) 100 iterations of CV were performed on MEIS RPKM regression dataset, with 1x layer not present, or present with x0.25, x0.5, or x0.75 dimensionality reduction with equal probability. If the 1x layer was present, the activation was linear or ReLU with equal probability. These results were then sub-sampled to obtain 30 sets for MEIS RPKM model with ReLU activation, which were used to train the serial model. For stability validation, every transfer and serial model also included a newly trained base RPKM model.

Table S1: Hyper-parameter ranges for MEIS model selection and final settings.

| Hyper-parameter | min | max | step | up-binding | down-binding | RPKM |
|---|---|---|---|---|---|---|
| 1-layer CNN | | | | | | |
| N. of filters | 32 | 1504 | 32 | 416 | 192 | 800 |
| Filter length | 8 | 29 | 1 | 19 | 27 | 15 |
| Dense (hidden) | 0 | 160 | 32 | 128 | 32 | 64 |
| Dropout | 0 | 0.9 | 0.1 | 0.7 | 0.8 | 0.7 |
| Input length | 200 | 2000 | 200 | 400 | 600 | 600 |
| Batch size | 16 | 512 | *2 | 256 | 16 | 256 |
| CV iterations | 200 | | - | Training time: | | |
| Early stop (epochs) | 10 | | - | 95s | 352s | 1297s |
| 1-layer - transfer | | | | | | |
| Dense (hidden) | 0 | 160 | 32 | 32 | 32 | |
| Dropout | 0 | 0.7 | 0.1 | 0.6 | 0.6 | |
| Batch size | 8 | 128 | *2 | 128 | 64 | |
| CV iterations | 30 | | - | Training time: | | |
| Early stop (epochs) | 10 | | - | 61s | 38s | |
| Deep, serial | | | | | | |
| Dense (hidden) | 0 | 160 | 32 | 128 | 64 | |
| Dropout | 0 | 0.7 | 0.1 | 0.3 | 0.3 | |
| Batch size | 8 | 128 | *2 | 64 | 16 | |
| CV iterations | 30 | | - | Training time: | | |
| Early stop (epochs) | 10 | | - | 37s | 56s | |
| Deep, parallel | | | | 3-task | 2-task | Deep |
| N. of filters (pooling) | 32 | 512 | 32 | 320 | 288 | 192 |
| Filter length (pooling) | 3 | 24 | 1 | 16 | 18 | 17 |
| Max-pooling rate | 2 | | - | 2 | 2 | 2 |
| N. of blocks (pooling) | 3 | | - | 3 | 3 | 3 |
| N. of filters (dilation) | 32 | 384 | 32 | 224 | 256 | 352 |
| Filter length (dilation) | 3 | 7 | 1 | 3 | 7 | 4 |
| Dilation rate | 4 | | - | 4 | 4 | 4 |
| N. of blocks (dilation) | 6 | 8 | 1 | 8 | 7 | 7[1] |
| Channel reduction | 0.25 | 0.75 | 0.25 | 0.25 | 0.5 | 0.25 |
| Dropout | 0 | 0.4 | 0.1 | 0.2 | 0.2 | 0.2 |
| RF expansion[2] | 1.0 | 2.0 | 0.1 | 1.2 | 2.0 | 1.2 |
| Batch size | 8 | 128 | *2 | 128 | 16 | 64 |
| Batch size (classification) | 4 | 44 | 4 | 28 | 40 | - |
| LRF[3] (classification) | 10 | 100 | 1 | 40 | 33 | - |
| CV iterations | 30 | | - | Training time: | | |
| Early stop (epochs) | 20 | | - | 7135s | 21672s | 2717s |

[1] Search range in model selection of RPKM model was set between 5 to 10 blocks.
[2] Multiplies receptive field to get input size.
[3] Learning rate factor, divides learning rate for classification tasks.

##### 5.2 Classification test performance

Table S2: Test set prediction performance for classification tasks (downbinding).

| Predicted \ Real | BA1 down | BA2 down | PBA down | Non-DB |
|---|---|---|---|---|
| **1-layer CNN** | | | | |
| BA1 down | 327 | 52 | 375 | 4627 |
| BA2 down | 17 | 412 | 170 | 3919 |
| PBA down | 80 | 71 | 2749 | 8430 |
| Non-DB | 41 | 75 | 301 | 8459 |
| Recall | 0.7 | 0.68 | 0.76 | 0.33 |
| PR-AUC | 0.13 | 0.16 | 0.38 | 0.93 |
| F1 avg. | 0.284 | | | |
| **1-layer CNN - transfer** | | | | |
| BA1 down | 325 | 32 | 269 | 3964 |
| BA2 down | 38 | 469 | 238 | 5741 |
| PBA down | 52 | 40 | 2643 | 5523 |
| Non-DB | 50 | 69 | 445 | 10207 |
| Recall | 0.7 | 0.77 | 0.74 | 0.4 |
| PR-AUC | 0.18 | 0.17 | 0.47 | 0.93 |
| F1 avg. | 0.318 | | | |
| **Deep model - direct classification** | | | | |
| BA1 down | 238 | 14 | 161 | 1837 |
| BA2 down | 71 | 441 | 506 | 5604 |
| PBA down | 19 | 11 | 957 | 1920 |
| Non-DB | 137 | 144 | 1971 | 16074 |
| Recall | 0.51 | 0.72 | 0.27 | 0.63 |
| PR-AUC | 0.12 | 0.16 | 0.27 | 0.9 |
| F1 avg. | 0.332 | | | |
| **Deep model - parallel, 2-task** | | | | |
| BA1 down | 298 | 35 | 79 | 2273 |
| BA2 down | 14 | 372 | 61 | 2372 |
| PBA down | 56 | 36 | 2845 | 6226 |
| Non-DB | 97 | 167 | 610 | 14564 |
| Recall | 0.64 | 0.61 | 0.79 | 0.57 |
| PR-AUC | 0.22 | 0.23 | 0.52 | 0.95 |
| F1 avg. | 0.391 | | | |
| **Deep model - parallel, 3-task** | | | | |
| BA1 down | 353 | 44 | 216 | 3650 |
| BA2 down | 9 | 379 | 108 | 2345 |
| PBA down | 13 | 5 | 2028 | 2196 |
| Non-DB | 90 | 182 | 1243 | 17244 |
| Recall | 0.76 | 0.62 | 0.56 | 0.68 |
| PR-AUC | 0.19 | 0.22 | 0.52 | 0.94 |
| F1 avg. | 0.417 | | | |

Continued from page 3.

| Deep model - serial | | | | |
|---|---|---|---|---|
| Predicted \ Real | BA1 down | BA2 down | PBA down | Non-DB |
| BA1 down | 410 | 75 | 327 | 6478 |
| BA2 down | 4 | 371 | 44 | 1743 |
| PBA down | 14 | 9 | 2394 | 3309 |
| Non-DB | 37 | 155 | 830 | 13905 |
| Recall | 0.88 | 0.61 | 0.67 | 0.55 |
| PR-AUC | 0.24 | 0.25 | 0.54 | 0.94 |
| F1 avg. | 0.394 | | | |

Table S3: Test set prediction performance for classification tasks (upbinding).

| Deep model - serial | | | | |
|---|---|---|---|---|
| Predicted \ Real | BA1 up | BA2 up | PBA up | Non-DB |
| BA1 up | 270 | 21 | 81 | 2970 |
| BA2 up | 193 | 648 | 264 | 7650 |
| PBA up | 79 | 60 | 2907 | 6061 |
| Non-DB | 108 | 44 | 377 | 8754 |
| Recall | 0.42 | 0.84 | 0.8 | 0.34 |
| PR-AUC | 0.09 | 0.25 | 0.51 | 0.93 |
| F1 avg. | 0.308 | | | |
| **Deep model - parallel, 3-task** | | | | |
| Predicted \ Real | BA1 up | BA2 up | PBA up | Non-DB |
| BA1 up | 284 | 67 | 109 | 3106 |
| BA2 up | 68 | 427 | 47 | 2148 |
| PBA up | 58 | 92 | 2710 | 5461 |
| Non-DB | 240 | 187 | 763 | 14720 |
| Recall | 0.44 | 0.55 | 0.75 | 0.58 |
| PR-AUC | 0.08 | 0.2 | 0.48 | 0.93 |
| F1 avg. | 0.387 | | | |

##### 5.3 Receptive field calculation

Receptive field is calculated as the maximum span in the input sequence, change in which can affect the activation of neurons prior to the global pooling layer, see algorithm 1.

```
Result: Receptive field sequence length (nt).
RF = 1;
stride = 1;
for Each layer do
    if Convolutional then
        RF += (kernel length - 1) * dilation rate * stride;
    end
    else if Pooling then
        RF += (pooling rate - 1) * stride;
        stride *= pooling rate;
    end
end
```

**Algorithm 1**: Receptive field calculation for multilayer CNN.

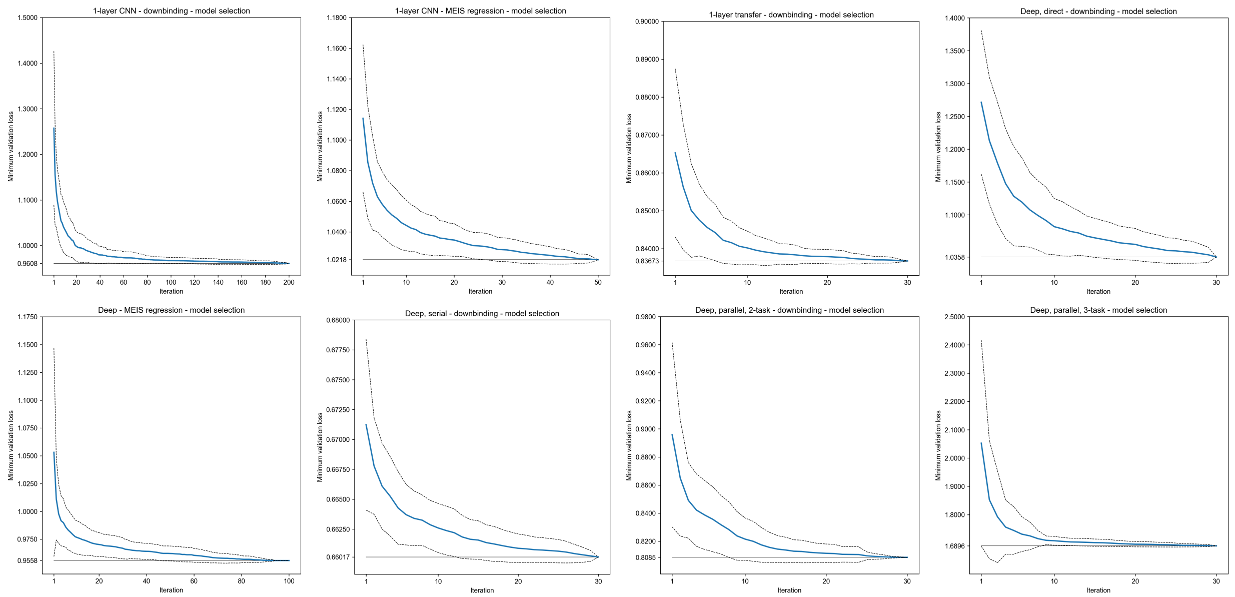


Figure S1: Lowest validation loss and standard deviation (dashed line) calculated over 100 permutations of hyper-parameter random search results.

Table S4: Reverse complement augmentation (downbinding PR-AUC)

| 1-layer CNN | | | | | |
|---|---|---|---|---|---|
| | BA1 down | BA2 down | PBA down | Non-DB | Avg. F1 |
| no RC | 0.11 | 0.12 | 0.31 | 0.91 | 0.25 |
| RC | 0.13 | 0.16 | 0.38 | 0.93 | 0.28 |
| Deep model - serial | | | | | |
| | BA1 down | BA2 down | PBA down | Non-DB | Avg. F1 |
| no RC | 0.24 | 0.25 | 0.54 | 0.94 | 0.36 |
| RC | 0.24 | 0.25 | 0.54 | 0.94 | 0.39 |
| Deep model - parallel, 3-task | | | | | |
| | BA1 down | BA2 down | PBA down | Non-DB | Avg. F1 |
| no RC | 0.19 | 0.21 | 0.51 | 0.93 | 0.38 |
| RC | 0.19 | 0.22 | 0.52 | 0.94 | 0.42 |

##### 5.4 Reverse complement augmentation

Reverse complement (a sequence of the complementary DNA strand, obtained by swapping C and G, T and A, and reversing order) is added to augment the the least frequent BA1-down class, doubling the example count. We only augment other down-binding classes if their example count is below the augmented BA1-down, resulting in partial augmentation of BA2-down. Other classes are not augmented. Augmentation is performed after splitting data into training and validation folds, to avoid leaking examples between them.

##### 5.5 K-mer counting methods

For Homer results we used *findMotifsGenome.pl* module to count in the differential regions, setting non-differential regions as background. 200nt input length was used with motif length $k$ from 5 to 12. HOXA2 motif is the most confident PWM in BA1-downbinding when counting *de novo* and appears as *(Pdx1(Homeobox)/Islet-Pdx1-ChIP-Seq(SRA008281)/Homer)* in known results. To annotate the regions *scanMotifGenomfeWide.pl* module was used with LOD=1 to find the motif genome-wide. Results were intersected with MEIS BA1-down regions and a single location with the highest LOD was selected in each region.

To obtain KSM/KMAC results we ran KMAC with MEIS BA1-down *fasta* file, using non-differential regions *fasta* as background. Motif length 5 to 12 was used. Similarly to Homer, Hox is the first cluster in the resulting prediction.

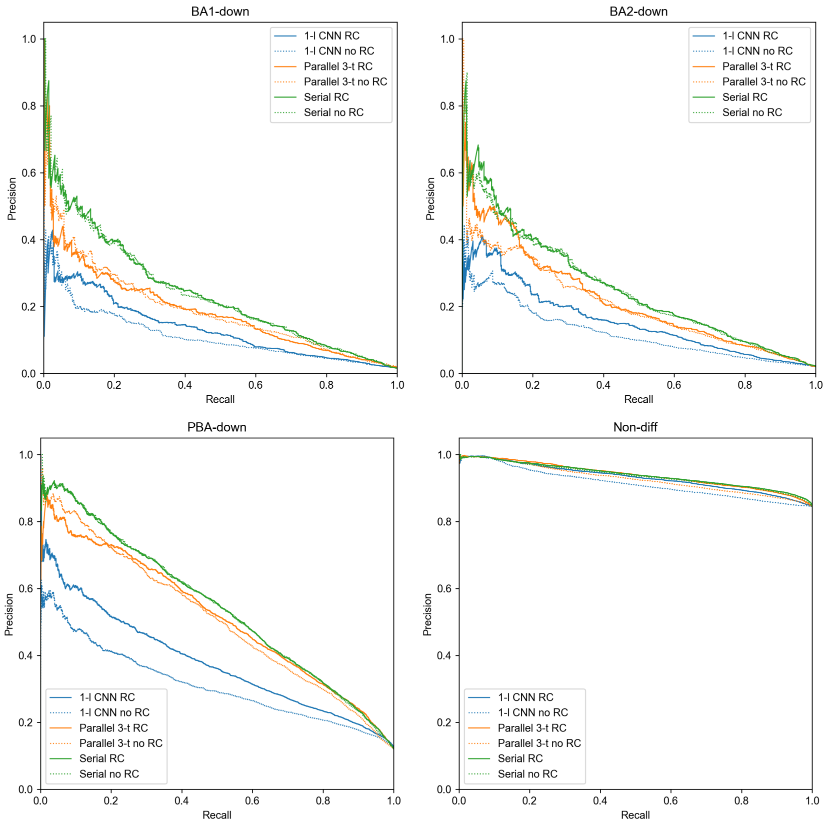


Figure S2: Precision-recall curves for model selection with and without RC augmentation. To increase class balance of BA1-down, only this class was fully augmented. BA2-down was partially augmented to match the count of augmented BA1-down. PBA and non-differential classes were not augmented.

We only considered k-mers in the first cluster and for each BA1-down region identified the location of k-mer with the lowest p-value.

##### 5.6 Mutagenesis

Mutagenesis scores are obtained by changing each of the bases in a region to its three alternatives and using the scoring function defined in DeepBind supplementary material [2]. For classification models attribution is obtained from class-specific activations preceding the softmax layer. Regions larger than 2000nt are trimmed to this size around the centre, for smaller regions mutations are evaluated only within the original span. If the region is smaller than the input size of the model, it is expanded and centred within the model input. For regions larger than model input we use a sliding approach, in which the attributed nucleotide is always centred in the input, except when the nucleotide is positioned closer to region start or end than half of model input width, in which case the input is not moved past the region span.

##### 5.7 Integrated gradients

Similarly to mutagenesis pre-softmax activations are used, regions are trimmed to a maximum of 2000nt and centred if smaller than model input. For regions larger than model input attribution is performed in strides, with overlap bigger or equal to 0.25 of model input width. Nucleotides with overlapping attribution have their scores averaged. When attributing using several references scores are combined by arithmetic mean. Summation-to-delta in Fig. S6 is calculated as $|S_{attr} - P_d| \,/\, |P_d|$, where $S_{attr}$ is the sum of attribution in a region, and $P_d$ is the difference in model prediction between the region and a reference. Values are calculated for 100 regions, 10 references each, and averaged.

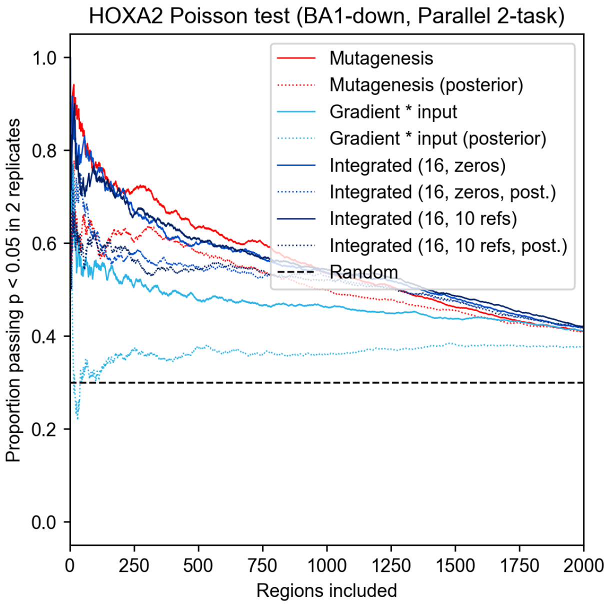


Figure S3: Attribution performance comparison when class posterior values (obtained after final softmax) are used, compared to using raw unnormalised logit values (default).

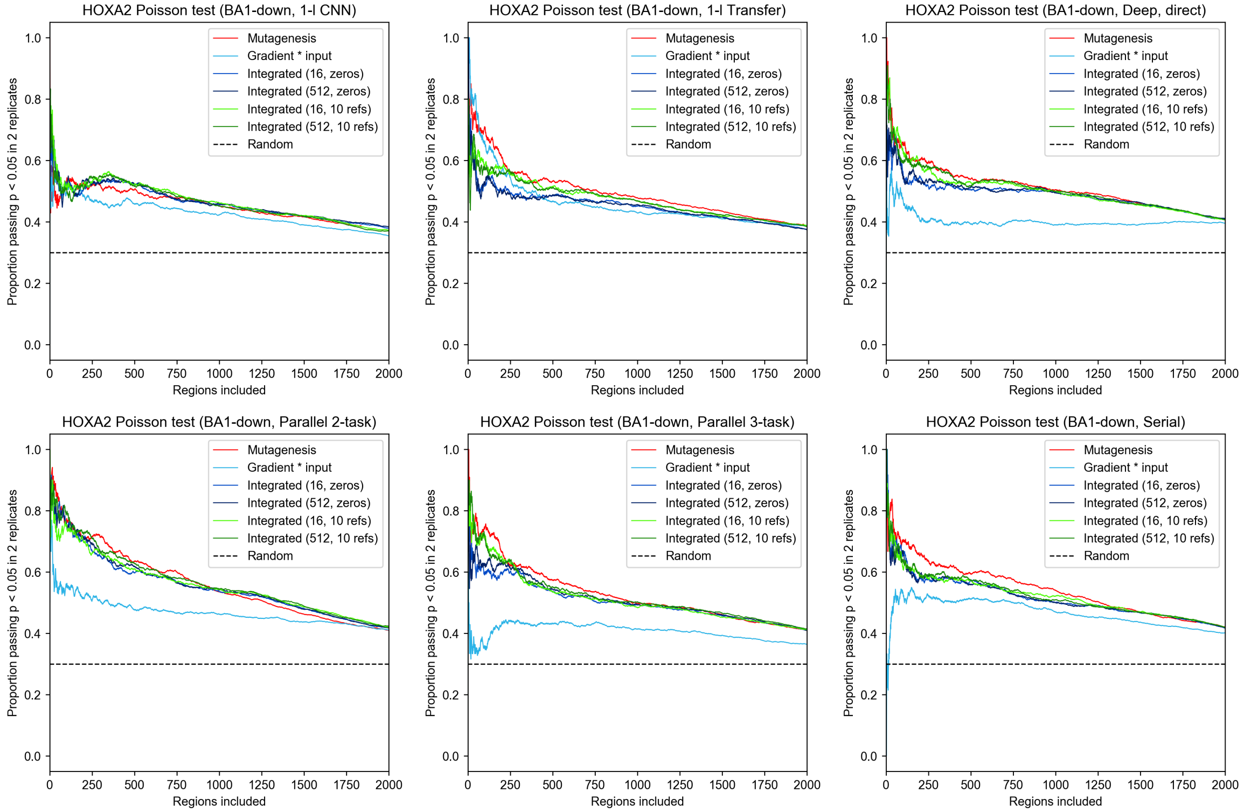


Figure S4: Attribution method comparison for BA1-downbinding for all deep learning models.

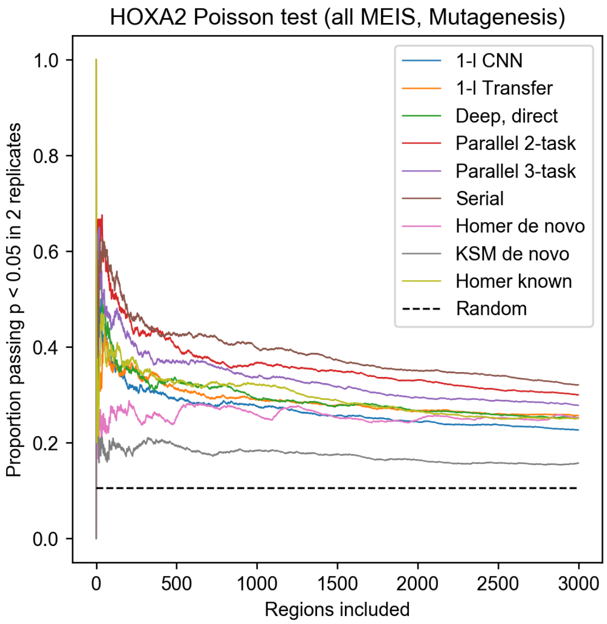


Figure S5: Model comparison for attribution performed in all MEIS-bound regions.

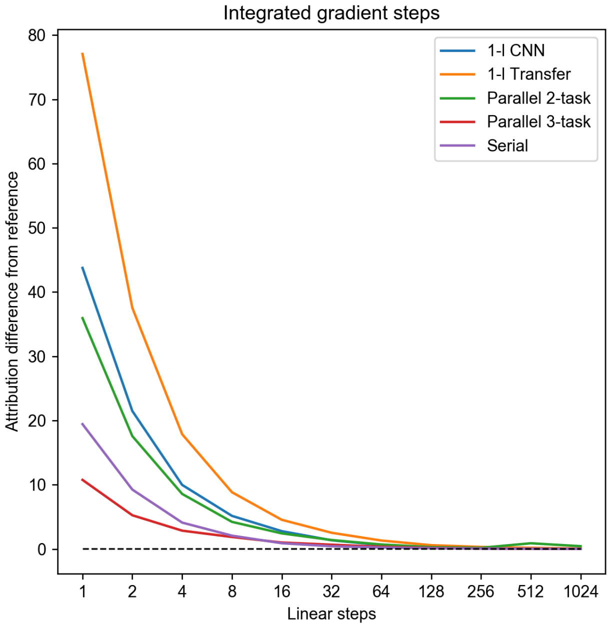


Figure S6: Relative difference between the absolute sum of attribution in a region and the difference in prediction between the region and a reference (summation-to-delta), as a function of increasing integration density of the gradient. Values above zero indicate over-complete explanation (manifesting as noisy attribution).

##### 5.8 Attribution peak calling

For all used attribution methods the results are equal in size to the one-hot input (region length * 4). Following attribution, one dimensional per-nucleotide scores are obtained by summing the 4 channels. For a given set of regions, strongest features are identified using a sliding window approach. A range of window lengths can be evaluated (11 to 25 in our tests, increasing by 2). To identify locations of strongest features of length $k$ in $r$ regions, based on one-dimensional attribution scores, we create a $r*k$ scores array and convolve it with a length-$k$ vector of values all equal to $1/k$, for all desired values of $k$, and keep the maximum value for each nucleotide. These scores are then sorted from strongest to weakest. To avoid the sliding window prioritising offsets which avoid strong negative scores, sorting is performed on absolute values of attribution first, after which only a fixed number of strongest features is kept (20000 for BA1-down regions only and 50000 for all MEIS regions). The remaining features are sorted again based on their original (non-absolute) scores and non-positive features are discarded (see Fig. S7 for ablation results). For Poisson and stability tests we find a strongest single feature per peak from the resulting list, if at least one feature is present in a peak, and select the 25nt region around its centre.

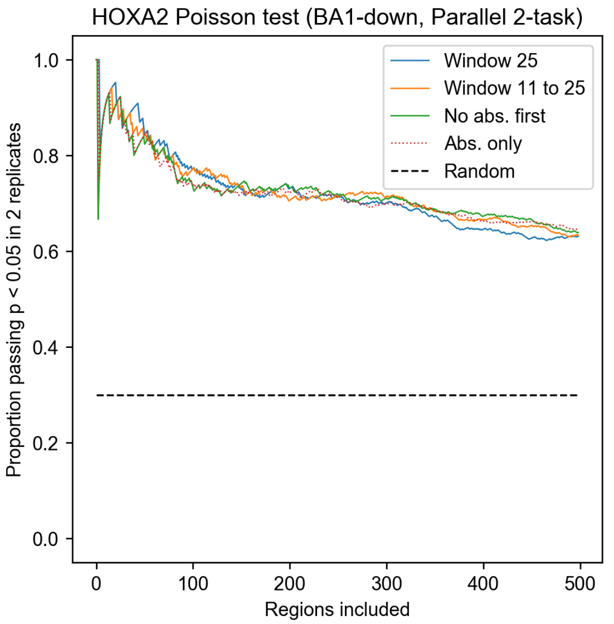


Figure S7: Ablation of attribution peak calling method performed in BA1-down regions. Window 25, or 11 to 25 (used in the paper), indicates the window size in which the attribution is summed around each nucleotide, keeping the maximum value. No abs. first indicates summing and ranking performed on the attribution values directly, without using absolute values first. Abs. only indicates using only absolute values of attribution.

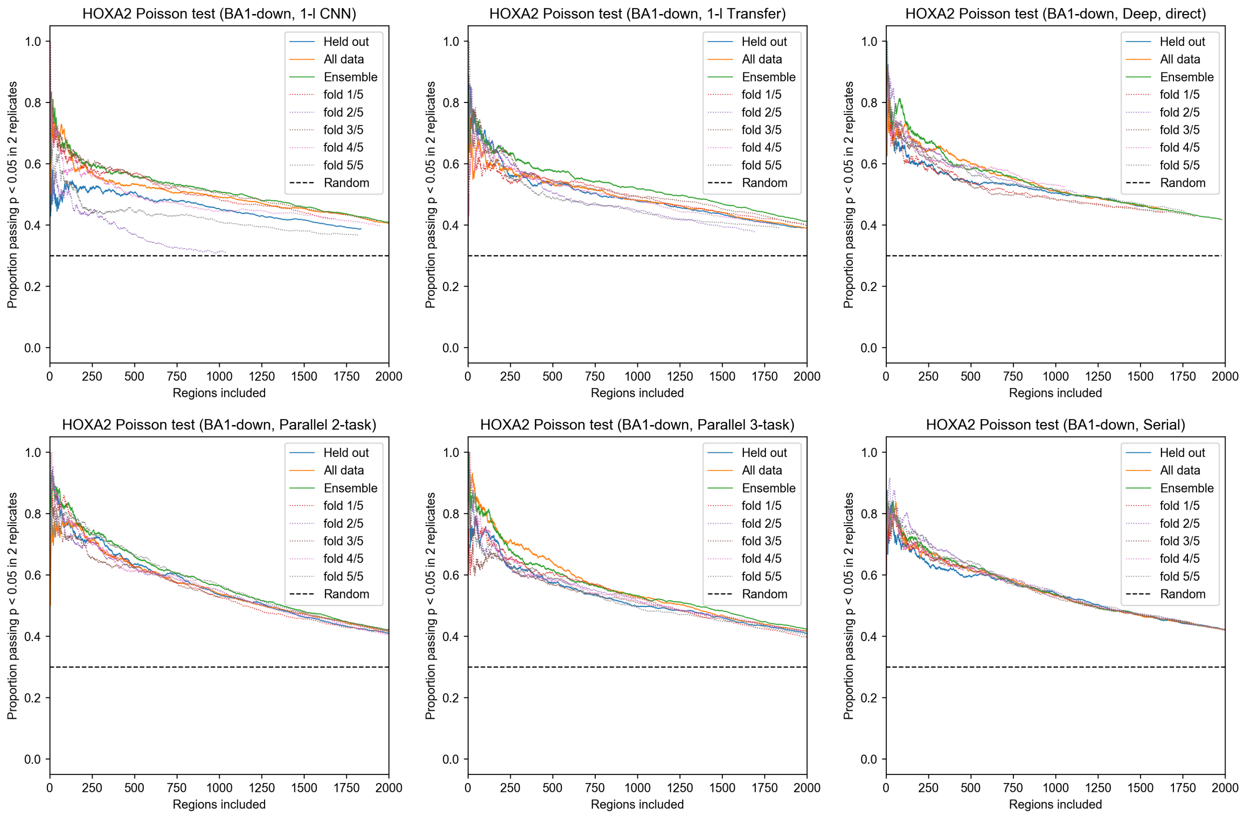


Figure S8: Overfitting effects in BA1-down mutagenesis attribution for all models. 5 models were trained holding out different folds of randomly shuffled data. Held out indicates each peak was attributed with model which held out the region during training. Ensemble indicates using mean attribution from all models. All data indicates using a single model, trained on all the data.

#### 5.9 Poisson test

Poisson test is performed on the identified feature locations using the approach in MACS2 [3], with 500nt window around each feature as suggested in GEM [4]. Test is performed separately on two HOXA2 BA2 ChIP-seq replicates and p<0.05 is required in both in order to pass. As in MACS2, we calculate:

$$\Lambda_{max} = max(\Lambda_{BG}, \Lambda_{500}, \Lambda_{1000}, \Lambda_{5000}, \Lambda_{10000}) \tag{1}$$

where $\Lambda_{BG}$ is the expected lambda value for a random 500nt region, and the remaining lambdas are calculated for the tested region in the corresponding sequence span. We perform total count normalisation across IP and Input ChIP-seq to make the values comparable, and use *ppois* command from the R package (3.5.1) to obtain the p-value: *ppois(Current window count,* $\Lambda_{max}$*, lower.tail = False).*

##### 5.10 Comparison with LS-GKM SVM

The performance of BA1-downbinding classification was compared to a gapped k-mer SVM method LS-GKM [5], which was evaluated with two types of kernels: wgkm and its radial basis expansion wgkmrbf. Since SVM models are binary classifiers, the predictive performance was compared with CNN models trained for binary task of classifying BA1-downbinding regions against the non-differential background. Precision-recall plot is shown in Fig. S9. Model selection for 1-layer CNN was performed for 50 iterations. CNN models outperform SVM in this task even without regularisation with regression data.

For attribution with SVM in-silico mutagenesis was used, as well as GkmExplain [6]. Results are shown in Fig. S10. For this comparison, non-binary CNN models were trained on the same classification data fold as the SVM. Therefore, unlike the main paper results, this attribution was performed using a single CNN model (fold 0) for the entire BA1-downbinding set. LS-GKM performs well in identifying the most confident features, and performs similarly to a shallow CNN, but is outperformed by the serial model as the number of included regions is increased. GkmExplain outperforms other methods in the top range, but the performance declines in the less confident regions to the advantage of mutagenesis. SVM with wgkm kernel outperforms wgkmrbf, and models trained on 600nt regions have better performance to those trained on 200nt. Fig. S11 illustrates the strongest features obtained from the SVM model by mutagenesis and GkmExplain. While the latter results in smoother features, it is perhaps detrimental to precise ranking.

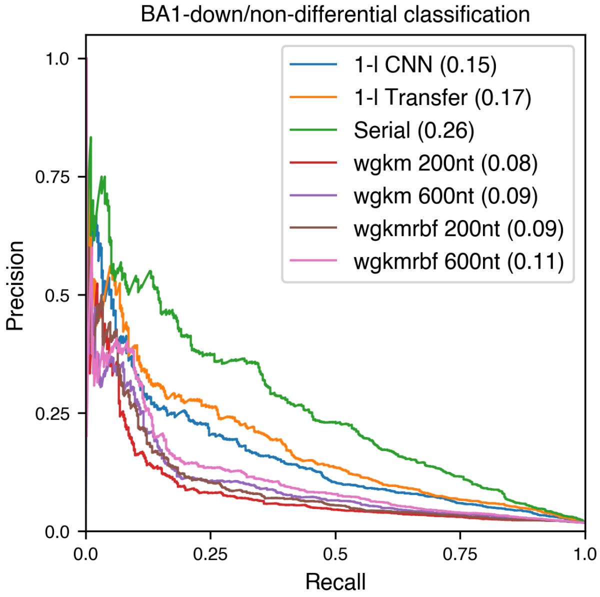


Figure S9: Comparison of CNN and LS-GKM SVM trained on a binary classification task. Transfer and serial models used regression data for regularisation. LS-GKM was trained with wgkm or wgkmrbf kernel and 200nt or 600nt region length. AUC shown in brackets.

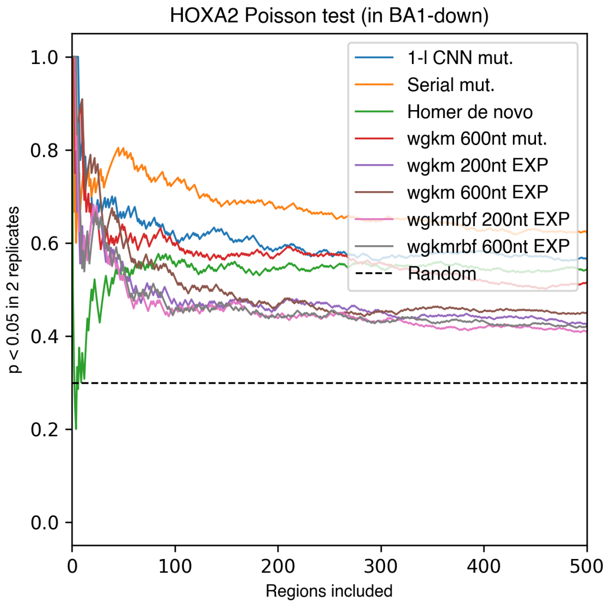


Figure S10: Comparison of attribution performance between CNN, Homer, and SVM in BA1-downbinding regions. CNNs used were the multitask models used throughout this chapter. LS-GKM models and Homer were trained on the binary classification tasks. For attribution with LS-GKM mutagenesis was used (mut.), as well as GkmExplain (EXP) with default settings.

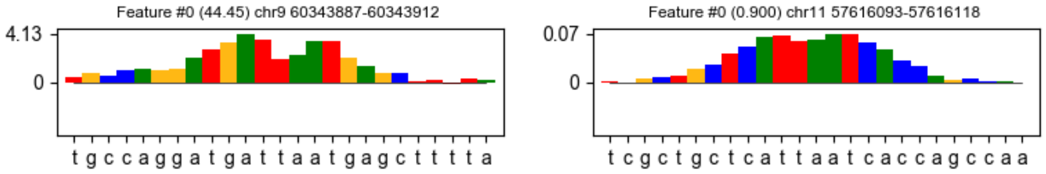


Figure S11: Strongest feature in BA1-downbinding regions identified by LS-GKM with wgkm kernel using in-silico mutagenesis (top) and GkmExplain (bottom).

##### 5.11 Time of model selection and attribution

Fig. S12 shows the total time it would take to perform model selection, including that of RPKM models, if a single GPU was used. In practise, model selection was performed in parallel on several nodes. Training times of models with known hyper-parameters are given in table S1. The majority of time spent on creating serial and transfer-like models is consumed by model selection of the base models, which can be reused for subsequent tasks. In training the shallower models, larger batch size can noticeably speed up the process due to better utilisation of GPU memory.

The comparison of model creation time between CNN, SVM and Homer was performed on a binary dataset, and the results are shown in table S5. Although deep learning models can take a long time to train, 1-layer CNNs for direct classification can be created from scratch in around an hour, which is much faster than LS-GKM on this dataset. However, comparing creation time between models is approximate, as the optimisation is performed with different hardware. In this work, neural networks were trained on a cluster consisting of Nvidia V100-SXM2 GPUs and Intel Skylake CPUs, with 8 CPU cores per GPU, benefitting from GPU acceleration in training and attribution. SVM and Homer were trained on Intel Broadwell CPUs with a total of 28 cores. Since these models do not require extensive hyper-parameter tuning, a single training time is reported.

Time of attribution per 1000 regions is given in table S6. CNN models were tested at their native input size, with batch size close to the limit imposed by GPU memory (16GB).

Table S5: Model creation time comparison, binary task (BA1-downbinding/non-differential, 600nt).

| Model | 1-layer CNN | | LS-GKM | Homer |
|---|---|---|---|---|
| | model selection | training | | |
| Time | 1h 35m 31s | 76s | 11h 27m 51s | 1h 54m 23s |

Table S6: Attribution time comparison, 1000 regions.

| Model | Input nt. | Method | Time |
|---|---|---|---|
| wgkmrbf | 600 | GkmExplain | 2h 32m 14s |
| 1-l CNN | 600 | Mutagenesis | 1m 47s |
| | | Grad.*Input | 117ms |
| | | Integ. 16 (zeros) | 2s |
| | | Integ. 512 (zeros) | 54s |
| | | Integ. 16 (10 refs) | 17s |
| | | Integ. 512 (10 refs) | 9m 18s |
| Deep CNN | 1000 | Mutagenesis | 12m 24s |
| | | Grad.*Input | 484ms |
| | | Integ. 16 (zeros) | 8s |
| | | Integ. 512 (zeros) | 4m 7s |
| | | Integ. 16 (10 refs) | 1m 18s |
| | | Integ. 512 (10 refs) | 41m 33s |

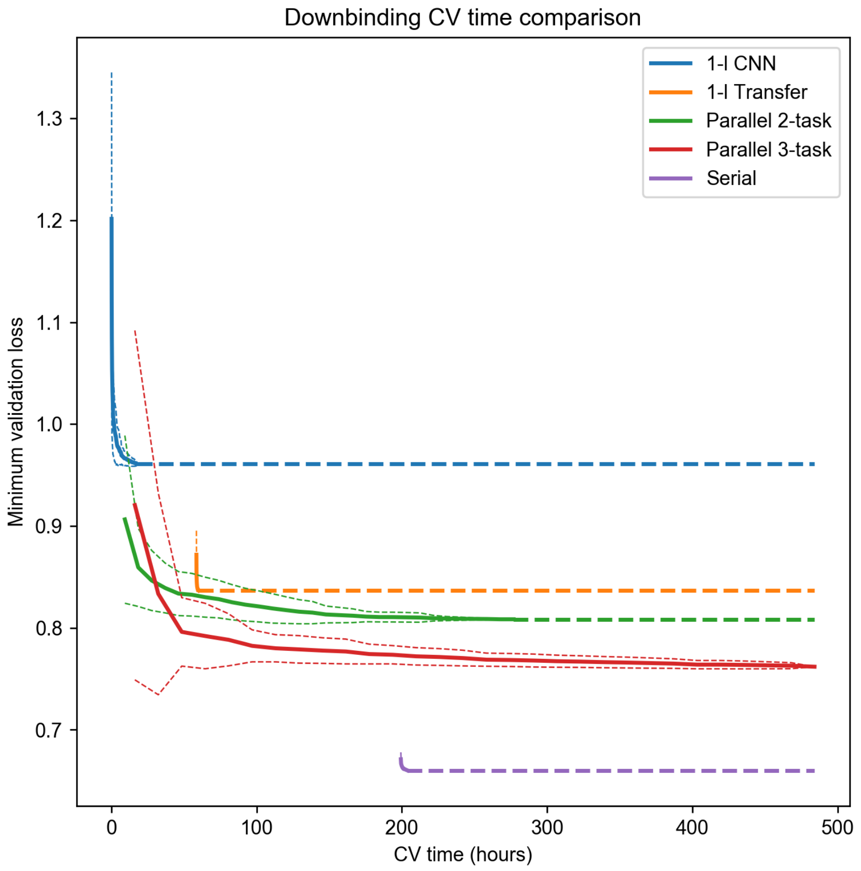


Figure S12: Validation loss on the down-binding task as a function of cross-validation time on a V100 GPU. Transfer and serial models are offset by a delay required to train their corresponding MEIS RPKM models (for 50 and 30 CV iterations respectively). Thin dashed lines represent standard deviation over 100 random permutations of the sampled hyper-parameter sets. Thick dashed lines represent final losses.

##### 5.12 MEIS up-binding features in PBA

HOX factors are known for their posterior prevalence, a phenomenon in which posterior HOX TFs (expressed in more posterior tissues) override the function of more anterior HOX. The parallel 3-task model was used to identify MEIS co-binding partners in PBA. Unlike in BA2, MEIS exhibits several co-binding partners in PBAs, examples of which are illustrated in Fig. S13.

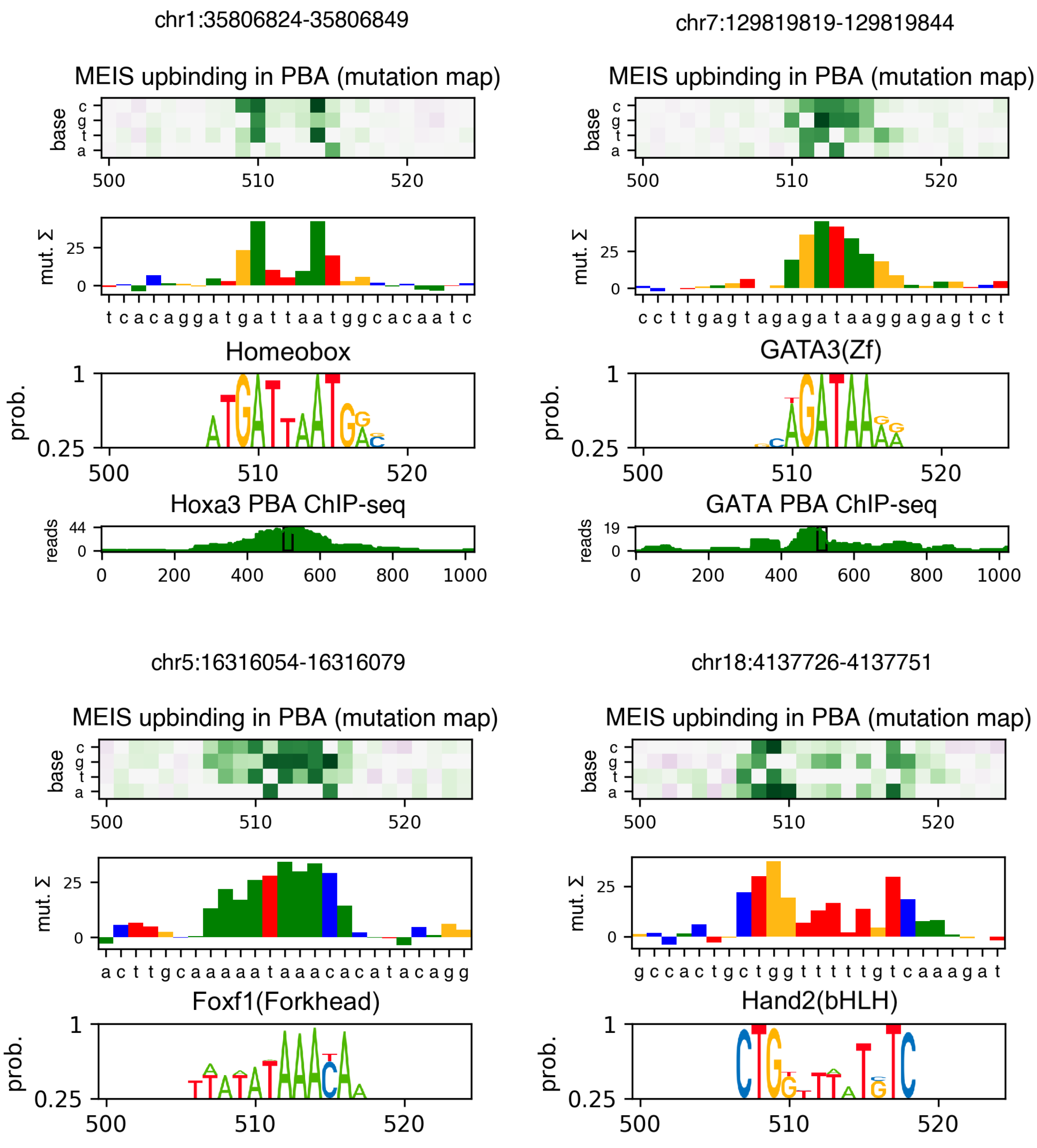


Figure S13: Example features of MEIS up-binding in PBA. MEIS exhibits several co-binding partners in this tissue. HOXA3 and GATA were validated by ChIP-seq. Forkhead (FOX) and HAND were validated by alignment with the known PWM annotated by Homer.

In order to cluster the features based on the underlying sequences, locations of highest ranked sites were identified in the PBA-upbinding regions using 25nt sliding window, and sorted by attribution sum. K-mer counting was performed in 100 thousand strongest features. Each feature was assigned the Homer annotation of a PWM with the highest sequence correlation. Fig. S14 shows maximum alignment of PBA-upbinding features with the PWMs.

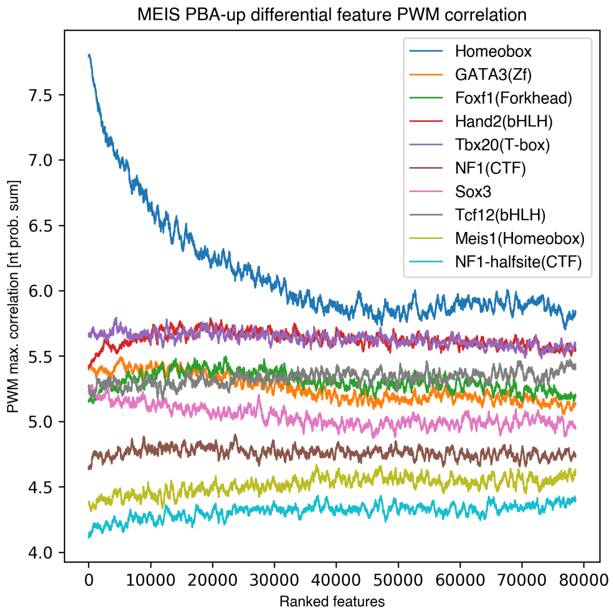


Figure S14: Correlation of strongest PBA-upbinding features with PWMs identified by Homer. Features were obtained from a 3-task parallel model using mutagenesis. Maximum cross-correlation with the underlying sequence and its reverse complement was calculated, and averaged across 500 ordered regions.

##### 5.13 MEF2D differential binding

MEF2D is a TF broadly expressed in a wide range of cells, regulating the development of tissues including neuronal, muscle, and retina [7]. It was shown to cooperate with CRX in mouse retina, allowing binding to non-specific motifs [8]. ChIP-seq data from mouse was analysed (retina: 3 replicates, cortical neurons: 4 experiments, myotubes: 2 replicates), and regions of up-binding and down-binding were identified. Sample counts are shown in table S8. For the baseline, 1-layer CNN classifiers were trained directly. For RPKM regression, a 1-layer CNN was trained for use with transfer-like models, and a deep CNN was trained for serial models, using between 1 and 5 dilation blocks in model selection (final setting $= 4$). Predictive performance of regression is shown in table S7, indicating that deep CNN increases performance over 1-layer model. Classification performance is shown in Fig. S15 and S16, table S9 and S10. Models regularised with regression result in significantly higher AUC than CNN trained directly. Deep, serial models exhibit higher PR-AUC than transfer-like models for most classes. Transfer-like CNN shows increased recall of the non-differential class, and results in highest F1 score in the down-binding task. The cortical neuron up-binding and retina down-binding classes show smaller variation in performance likely due to low example count and weak motif enrichment (top motif p-values identified by Homer are 1e-18 and 1e-19 respectively in those classes, counting against shuffled background).

Table S7: MEF2D cross-replicate and regression test correlation.

| | | Cross-replicate | | 1-l CNN | | Deep CNN | |
|---|---|---|---|---|---|---|---|
| Tissue | Replicate | R | Rho | R | Rho | R | Rho |
| Cortical neurons | 1 | - | - | 0.603 | 0.625 | 0.627 | 0.651 |
| Cortical neurons | 1 | - | - | 0.516 | 0.513 | 0.564 | 0.562 |
| Cortical neurons | 1 | - | - | 0.542 | 0.547 | 0.583 | 0.585 |
| Cortical neurons | 1 | - | - | 0.385 | 0.364 | 0.386 | 0.366 |
| Retina | 1 | 0.493 | 0.508 | 0.408 | 0.388 | 0.418 | 0.398 |
| Retina | 2 | 0.461 | 0.458 | 0.314 | 0.295 | 0.328 | 0.313 |
| Retina | 3 | 0.499 | 0.508 | 0.381 | 0.365 | 0.392 | 0.382 |
| Myotubes | 1 | 0.708 | 0.671 | 0.667 | 0.597 | 0.712 | 0.649 |
| Myotubes | 2 | 0.708 | 0.671 | 0.59 | 0.564 | 0.657 | 0.645 |

Table S8: Differential labelling of MEF2D-bound regions.

| Type | Count | Avg. length (nt) |
|---|---|---|
| Increased binding | | |
| Cortical neurons | 414 | 268.7 |
| Retina | 2344 | 510.7 |
| Myotubes | 3133 | 487.2 |
| Decreased binding | | |
| Cortical neurons | 452 | 629.0 |
| Retina | 604 | 395.5 |
| Myotubes | 2358 | 423.2 |
| Non-differential | 80131 | 306.3 |
| All MEF2D | 96529 | 323.6 |

Table S9: Test set prediction performance for MEF2D classification (upbinding).

| **1-layer CNN** | | | | |
|---|---|---|---|---|
| | *Real* | | | |
| *Predicted* | Cortical neurons up | Retina up | Myotubes up | Non-DB |
| Cortical neurons up | 56 | 50 | 106 | 3743 |
| Retina up | 8 | 344 | 144 | 4621 |
| Myotubes up | 9 | 54 | 351 | 5375 |
| Non-DB | 3 | 7 | 39 | 2285 |
| Recall | 0.737 | 0.756 | 0.548 | 0.143 |
| PR-AUC | 0.056 | 0.158 | 0.071 | 0.962 |
| F1 avg. | 0.127 | | | |
| **1-layer CNN - transfer** | | | | |
| | *Real* | | | |
| *Predicted* | Cortical neurons up | Retina up | Myotubes up | Non-DB |
| Cortical neurons up | 48 | 34 | 44 | 1964 |
| Retina up | 5 | 332 | 68 | 1780 |
| Myotubes up | 10 | 51 | 420 | 4827 |
| Non-DB | 13 | 38 | 108 | 7453 |
| Recall | 0.632 | 0.73 | 0.656 | 0.465 |
| PR-AUC | 0.064 | 0.313 | 0.104 | 0.977 |
| F1 avg. | 0.267 | | | |
| **Deep CNN - serial** | | | | |
| | *Real* | | | |
| *Predicted* | Cortical neurons up | Retina up | Myotubes up | Non-DB |
| Cortical neurons up | 60 | 33 | 50 | 3219 |
| Retina up | 1 | 363 | 40 | 1229 |
| Myotubes up | 10 | 42 | 439 | 5140 |
| Non-DB | 5 | 17 | 111 | 6436 |
| Recall | 0.789 | 0.798 | 0.686 | 0.402 |
| PR-AUC | 0.062 | 0.432 | 0.152 | 0.974 |
| F1 avg. | 0.273 | | | |

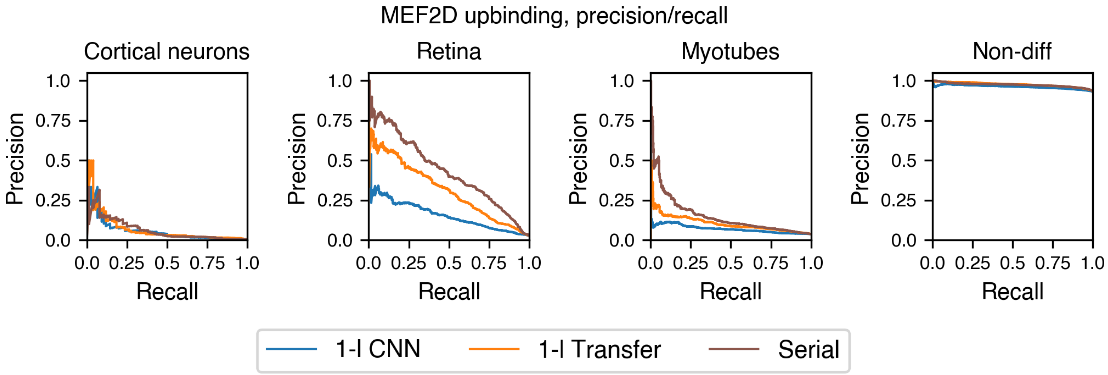


Figure S15: Test set precision/recall curves for the MEF2D differential upbinding task. 1-layer CNN was trained directly on the classification data. Transfer and serial models used regression data for regularisation.

Table S10: Test set prediction performance for MEF2D classification (downbinding).

| **1-layer CNN** | | | | |
|---|---|---|---|---|
| | *Real* | | | |
| *Predicted* | Cortical neurons down | Retina down | Myotubes down | Non-DB |
| Cortical neurons down | 51 | 19 | 78 | 2552 |
| Retina down | 13 | 66 | 60 | 5891 |
| Myotubes down | 5 | 2 | 240 | 891 |
| Non-DB | 17 | 33 | 99 | 6690 |
| Recall | 0.593 | 0.55 | 0.503 | 0.417 |
| PR-AUC | 0.068 | 0.011 | 0.322 | 0.983 |
| F1 avg. | 0.235 | | | |
| **1-layer CNN - transfer** | | | | |
| | *Real* | | | |
| *Predicted* | Cortical neurons down | Retina down | Myotubes down | Non-DB |
| Cortical neurons down | 59 | 13 | 35 | 1419 |
| Retina down | 10 | 74 | 22 | 5148 |
| Myotubes down | 7 | 4 | 353 | 1546 |
| Non-DB | 10 | 29 | 67 | 7911 |
| Recall | 0.686 | 0.617 | 0.74 | 0.494 |
| PR-AUC | 0.14 | 0.018 | 0.417 | 0.987 |
| F1 avg. | 0.264 | | | |
| **Deep CNN - serial** | | | | |
| | *Real* | | | |
| *Predicted* | Cortical neurons down | Retina down | Myotubes down | Non-DB |
| Cortical neurons down | 77 | 18 | 38 | 2044 |
| Retina down | 3 | 72 | 17 | 6803 |
| Myotubes down | 3 | 1 | 388 | 1667 |
| Non-DB | 3 | 29 | 34 | 5510 |
| Recall | 0.895 | 0.6 | 0.813 | 0.344 |
| PR-AUC | 0.175 | 0.03 | 0.475 | 0.989 |
| F1 avg. | 0.223 | | | |

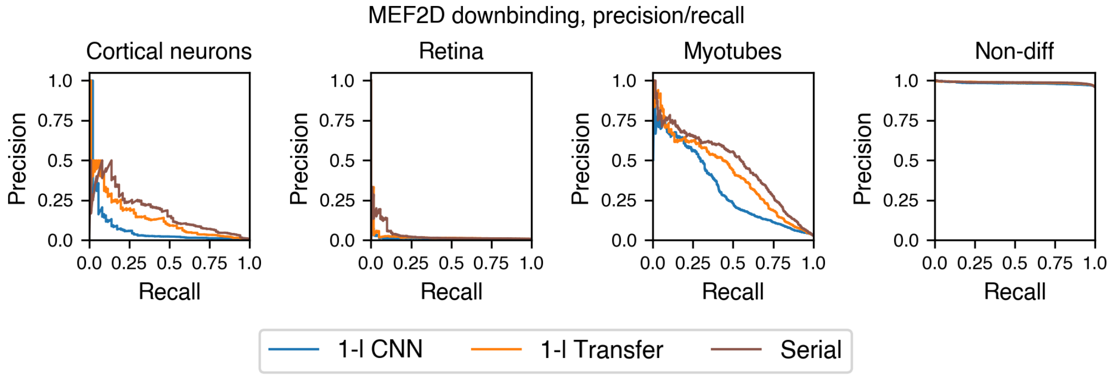


Figure S16: Test set precision/recall curves for the MEF2D differential downbinding task. 1-layer CNN was trained directly on the classification data. Transfer and serial models used regression data for regularisation.

Both transfer-like and serial models were subsequently used to obtain features of each class. In particular, Fig. S17 illustrates two example regions where differential features of MEF2D up-binding in retina consist of neighbouring MADS-domain and CRX binding sites. Example features for all classes are shown in Fig. S18. MADS binding sites appear as features for all up-binding classes, suggesting that co-factors and the surrounding sequence are important for specific differentiation. The models uncover known co-factors, such as CRX in retina [8] and MYOD in mytoubes [7]. Other possible co-factors were also identified, such as BATF in myotubes and cortical neurons (or other AP-1 factor, previously reported to interact with MEF2C by [9]), OTX in myotubes down-binding (a CRX-like factor), and IRF4 in cortical neurons up-binding (reported to interact with MEF2 in B-cells, [10]).

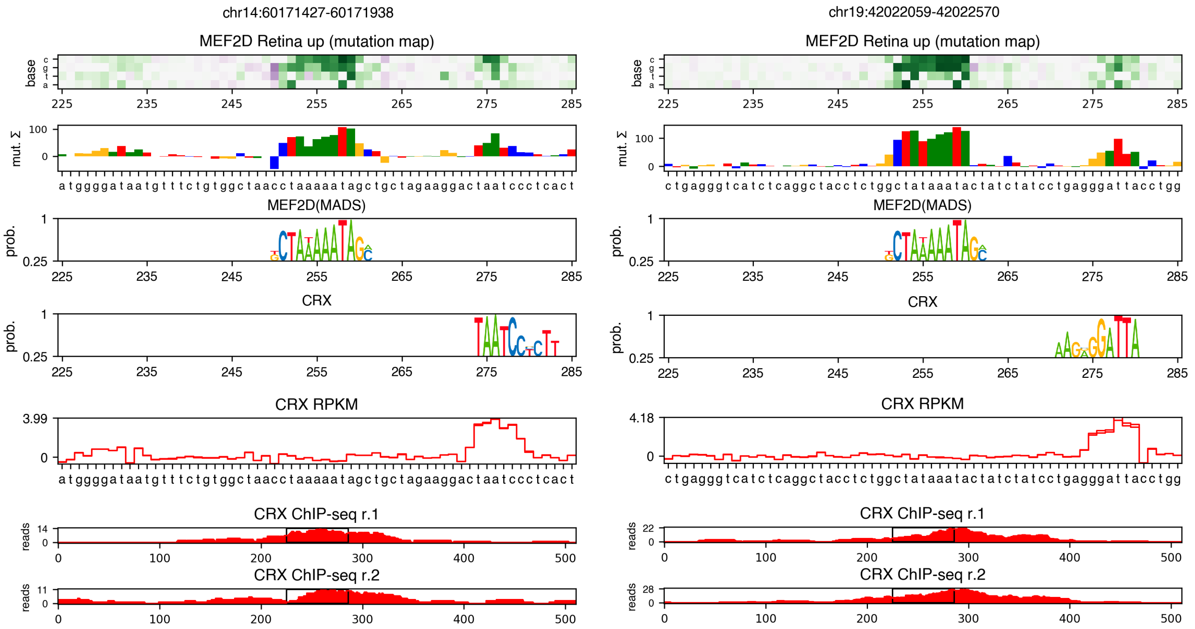


Figure S17: Example regions of MEF2D differential up-binding in mouse retina compared to myotube and cortical neuron tissues. Differential features were identified by a serial model using mutagenesis. PWMs for MEF2D and CRX were aligned to the underlying sequence and shown in the position of highest cross-correlation. CRX RPKM features are shown below, as well as CRX ChIP-seq profile in expanded regions. RPKM features were obtained from a model trained on CRX ChIP-seq data (not used to train the differential model).

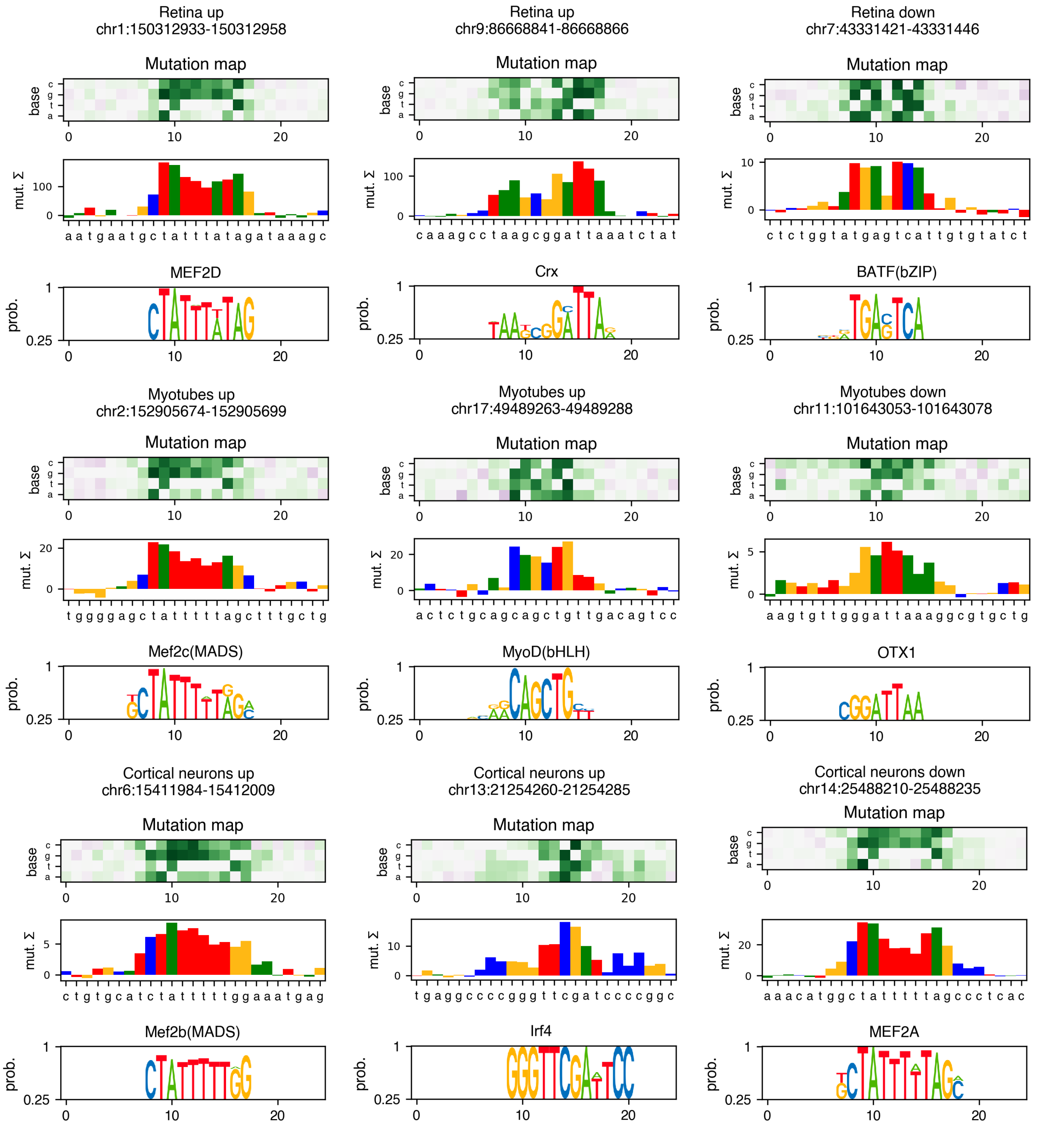


Figure S18: Example features of MEF2D differential binding in mouse retina, myotube, and cortical neuron tissues, identified by transfer-like or serial models using mutagenesis. Homer annotation of the enriched k-mers is shown below the differential tracks.

[5] Dongwon Lee. LS-GKM: A new gkm-SVM for large-scale datasets. *Bioinformatics*, 2016.

[6] Avanti Shrikumar, Eva Prakash, and Anshul Kundaje. GkmExplain: Fast and accurate interpretation of nonlinear gapped k-mer SVMs. In *Bioinformatics*, 2019.

[7] Matthew J. Potthoff and Eric N. Olson. MEF2: A central regulator of diverse developmental programs, 2007.

[8] Milena M. Andzelm, Timothy J. Cherry, David A. Harmin, Annabel C. Boeke, Charlotte Lee, Martin Hemberg, Basil Pawlyk, Athar N. Malik, Steven W. Flavell, Michael A. Sandberg, Elio Raviola, and Michael E. Greenberg. MEF2D drives photoreceptor development through a genome-wide competition for tissue-specific enhancers. *Neuron*, 2015.

[9] Teruyo Nakatani and Nicola C. Partridge. MEF2C interacts with c-FOS in PTH-stimulated Mmp13 gene expression in osteoblastic cells. *Endocrinology*, 2017.

[10] Julia Herglotz, Ludmilla Unrau, Friderike Hauschildt, Meike Fischer, Neele Kriebitzsch, Malik Alawi, Daniela Indenbirken, Michael Spohn, Ursula Müller, Marion Ziegler, Wolfgang Schuh, Hans Martin Jäck, and Carol Stocking. Essential control of early B-cell development by Mef2 transcription factors. *Blood*, 2016.
